# The Fitness Cost of Therapeutic Resistance in Cancer: A Systematic Review

**DOI:** 10.64898/2025.12.29.696883

**Authors:** Bailey Kane, Lauren Mestas, Madds Garza, Tiara Soesilo, Meghan Hufford, Harley Richker, Carlo Maley

**Affiliations:** Arizona Cancer Evolution Center, Arizona State University, Tempe, Arizona, USA 85287-5301; Biodesign Center for Biocomputing, Security and Society, Arizona State University, Tempe, Arizona 85287; School of Life Sciences, Arizona State University, Tempe, Arizona, USA 85281; School of Molecular Sciences, Arizona State University, Tempe, Arizona, USA 85281; Center for Evolution and Medicine, Arizona State University, Tempe, Arizona, 85281

## Abstract

Emergence of therapeutic resistance is a critical clinical challenge in cancer treatment, contributing to treatment failure, disease relapse, and patient mortality. Prior to treatment initiation, therapy-resistant clones typically make up such a small proportion of the neoplasm that they can be hard to detect and only expand under the selective pressure of therapy. This suggests that resistant clones incur a fitness cost for resistance to therapy. Whether therapeutic resistance incurs a fitness cost has important clinical implications, including the potential to leverage competition with sensitive cells to prevent the expansion of the resistant population. However, the fitness compromises associated with treatment resistance in cancer are not well-understood at present. We conducted a systematic review for peer-reviewed papers that fulfilled the following selection criteria: (i) experiments depict direct competition, (ii) between therapy-resistant and therapy-sensitive clones, (iii) in an untreated environment. We found 49 experiments that matched those criteria, of which approximately two-thirds found a fitness cost to resistance in a competitive environment. Of all pooled features, we found that the resistance characteristic was most significantly associated with whether resistant clones exhibited a fitness advantage in competition (p=0.0154). Further, we identify complex ecological interactions that may influence the behavior of the cancer cell population in the untreated environment. Predicting which resistance characteristics can be exploited therapeutically and identifying potential methods of modulating the costliness of the resistant phenotype may be critical to future improvements in cancer therapy.

**Lay Summary:** When cancer cells evolve to no longer respond to a cancer therapy, do they pay a cost for that invulnerability? We reviewed experiments where cancer cells resistant to a therapy were pitted against cancer cells that were still sensitive to the therapy. Approximately ⅔ of the experiments showed that the sensitive cells could out-compete the resistant cells, implying that the latter were paying some cost for their ability to resist the therapy.

## Overview

### The Clinical Challenge of Resistance in Cancer

Despite major strides in interventional strategies for cancer management, therapeutic resistance represents a major clinical challenge driving treatment failure, disease recurrence, and poor patient outcomes [1]. Standard-of-care approaches are typically administered using a fixed dosing strategy designed to eliminate as many cancer cells as possible within a short window of time. Typically, this dose is determined as the concentration that induces the highest cytotoxicity for the cancer without causing intolerable side effects for the patient: the Maximum Tolerated Dose (MTD). MTD treatment often induces appreciable tumor response initially, but can inadvertently enable the selective expansion of therapy-resistant clones that persist through treatment [2]. Thus, the tumor cell population may become refractory to further intervention, leading to malignant recurrence and therapeutic failure. Therapeutic resistance is frequently cited as a contributor to approximately 90% of cancer-related deaths [3]. However, the precise origin of this statistic is unclear, suggesting it is a heuristic rather than a precise measurement. Nevertheless, the emergence of resistance substantially limits treatment options and remains a barrier to long-term cancer control. Addressing therapeutic resistance is therefore critical both to patient care and the development of new-generation therapeutics.

Whether or not therapeutic resistance imposes a fitness cost on cancer cells has important clinical implications. If resistance is costly, then resistant cell populations will tend to be rare at diagnosis and thus hard to detect, which is a challenge for accurate prediction of the efficacy of a therapy. However, if resistance is costly, e.g., if the resistant clone grows slowly, it may take longer for the resistant clone to expand and for the therapy to fail than if resistance is not costly. Cancer drugs that require more costly forms of resistance should improve time to progression and overall survival compared to cancer drugs for which resistance phenotypes provide a fitness advantage even before the drug is applied. Further, we might use drug holidays or drug dose reductions to allow sensitive cells to out-compete resistant cells and thereby control the cancer – a strategy known as Adaptive Therapy (AT) [4]. That strategy, and many models of the evolution of therapeutic resistance [5,6], assume that there is some fitness cost to therapeutic resistance. Is that assumption justified?

### Evolution and Mechanisms of Resistance

Therapeutic resistance is a phenomenon observed across systems [7,8]. Within the scope of this review, we define therapeutic resistance as the ability of a tumor cell to evade or withstand cytotoxic treatment. Primary resistance notwithstanding (Explanatory Box 1), tumors may acquire resistance following a period of therapeutic pressure through two mechanisms: (i) selection for pre-existing resistant clones within a heterogeneous, treatment-naive tumor population, or (ii) the emergence of *de novo* resistant clones from sensitive parental lines [5,6]. Although conceptually distinct, these mechanisms are not mutually exclusive. In addition, resistance characteristics can arise through genetic or non-genetic changes, including reversible phenotypic alterations (phenotypic plasticity), which enable the persistence of drug-tolerant cell states [6,9]. Importantly, both genetic and non-genetic mechanisms can be heritable and therefore relevant to evolution by natural selection [10,11].

## Explanatory Box 1. Primary and Acquired Resistance

Primary resistance is a flexible term often used to describe both tumors and the individual cells that comprise them, though a tumor may appear to be resistant even if it initially responds to the drug but regrows before the therapeutic effects are measured. For the purpose of this review, we define primary resistance as the inherent tolerance of a cancer to a particular treatment, which presents clinically as nonresponse to treatment. Acquired resistance, on the other hand, is the emergence of resistance over the course of treatment that presents clinically as a gradual inefficacy of therapy following an initial response [12–14]. The confusion arises as such: acquired resistance of a cancer can result from selection on cells bearing preexisting resistance characteristics, which may well be considered primary resistance by some definitions. To avoid this mischaracterization, the terms primary and acquired resistance will refer only to entire tumors, and we will discuss the evolutionary mechanisms by which cancer cells possessing resistant phenotypes are driven to a majority.

**Figure 1.**
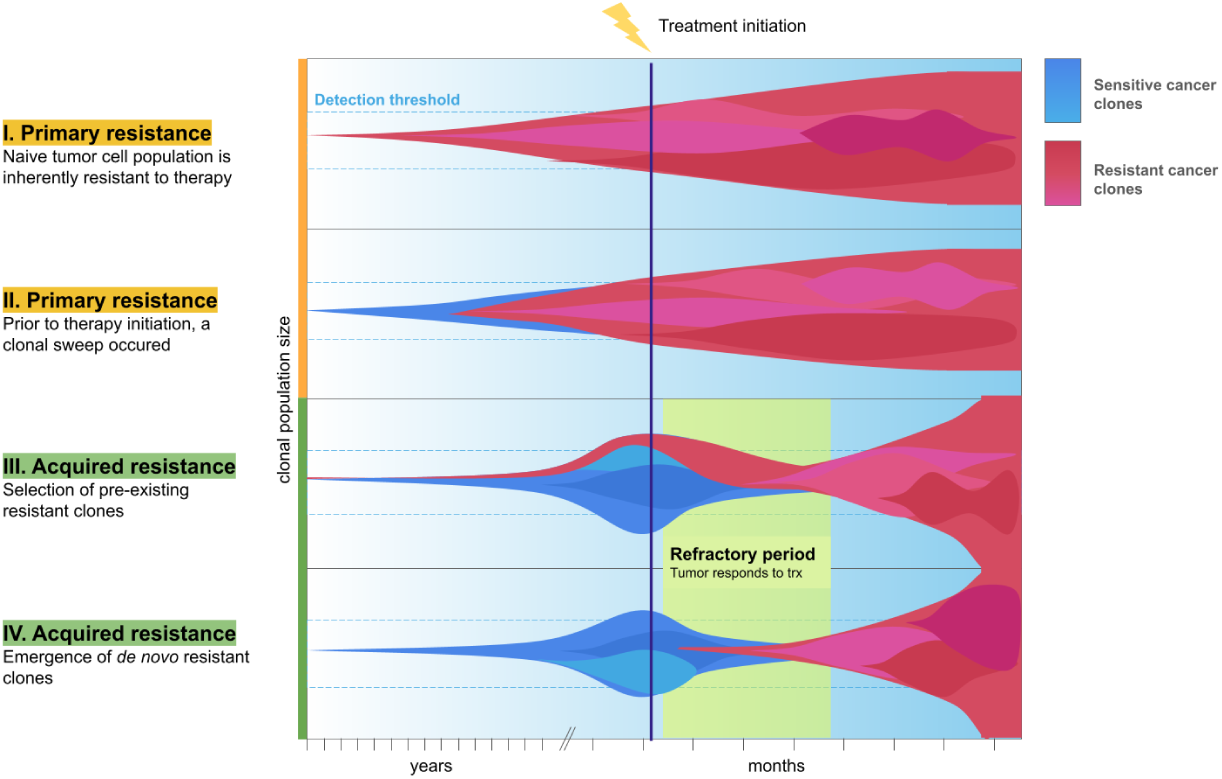
Clonal Expansion of Resistant Lineages. **Fig. 1** These fish plots illustrate primary and acquired resistance mechanisms. I. Primary resistance can result from a selective sweep of a resistant clone with high fitness even in the absence of therapy. II. Primary resistance can also occur as the therapy-naive tumor cell population is inherently nonresponsive to a treatment. III. Acquired resistance can result from selection of preexisting resistant mutants that comprise a small fraction of the therapy-naive population. IV. Therapeutic pressure can induce resistance acquisition by causing and/or promoting selection of de novo resistant phenotypes. Vertical axis: relative size of each clonal population in a tumor. Horizontal axis: time units. Note that the long period of neoplastic progression is compressed into units of years prior to therapy.

Combined with distinct evolutionary histories, the interaction of genetic and non-genetic factors generates various scenarios that explain the persistence of a resistant residual population following treatment. In the traditional view of Darwinian selection, pre-existing resistant clones in a heterogeneous population are capable of withstanding the therapy, producing therapy-tolerant offspring that clonally expand and lead to population-level resistance [15]. In a view of Lamarckian induction, however, a population of cancer cells may contain no pre-existing resistant individuals; instead, the resistant phenotype emerges as a result of gradual phenotypic habituation to therapy [16,17]. In a third scenario of dynamic fluctuation, plastic cells may transiently express a resistant phenotype and persist through treatment (called “persister cells”), but may variably switch phenotypes and shed the putative high-cost resistance program upon cessation of treatment [18,19]. A portion of these cells may enter a fixed resistant state [20]. Each model addresses how a population survives the evolutionary bottleneck imposed by therapy [14], but they are not mutually exclusive; tracing the evolutionary history of cancers reveals that multiple programs may be employed over the course of resistance acquisition [21].

Cancer cells employ various resistance strategies that enable persistence through treatment. Attempts to categorize mechanisms of resistance to cancer therapies have been numerous, though these frameworks often fail to be comprehensive, exhibit definitional redundancies, or conflate proximal mechanisms with regulatory mediators. A framework by Cree and Charlton identifies six distinct strategies for treatment evasion or compensation: alteration of drug targets, drug influx and efflux, drug metabolism, reduced apoptosis, altered proliferation, and increased DNA damage repair [22]. We expand this framework to accommodate insights from Longley and Johnson and include alternative or bypass signaling pathways in our review [3]. Gottesman et al. identify three major categories for cell-level mechanisms: Pharmacologic challenges, therapeutic target alterations, and cell survival adaptations (Figure 2) [23].

**Figure 2.**
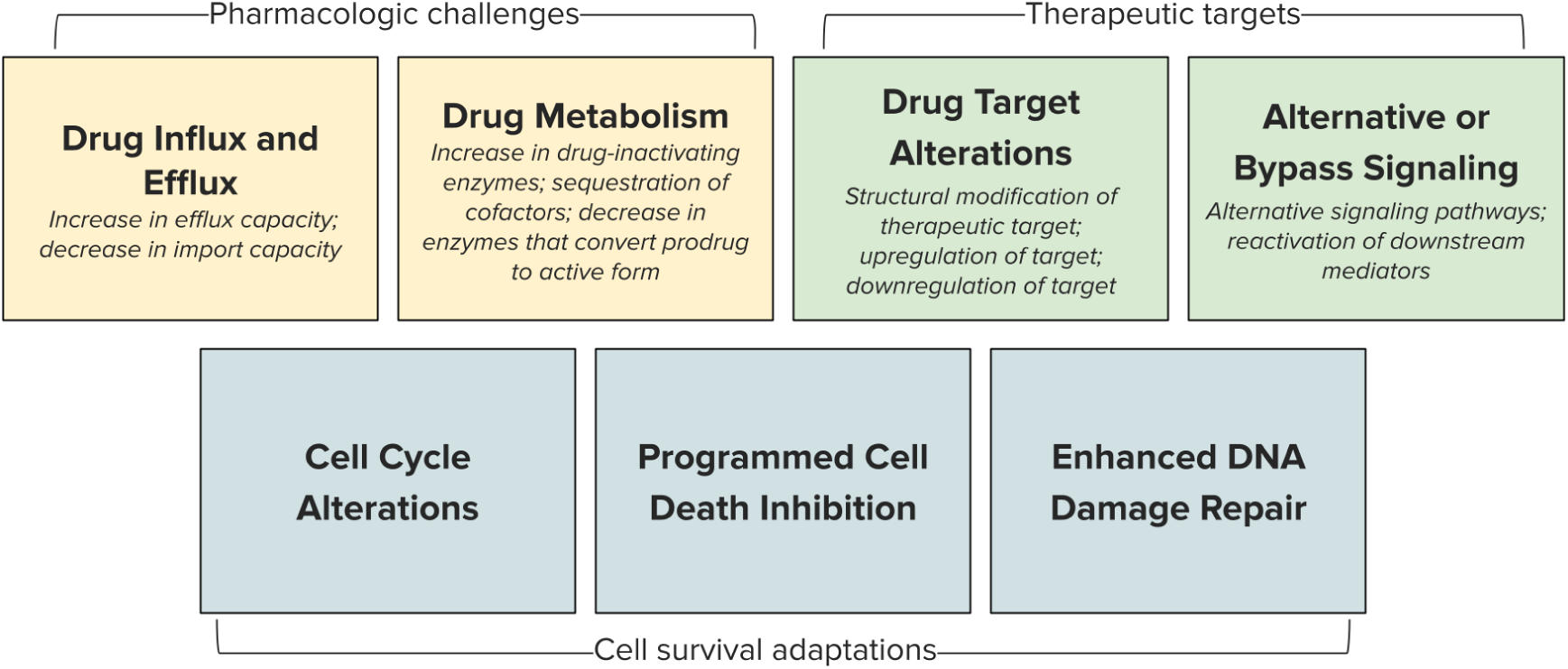
Cellular Mechanisms of Resistance. **Fig. 2** We recognize three general categories of mechanisms by which cells develop resistance to therapeutic agents. Pharmacologic challenges include drug delivery obstacles via efflux and influx and changes to drug metabolism. Cells may evade therapy through target alterations, including modifying the target’s structure or its abundance, or by relying on alternative signaling pathways. Cells may also undergo survival adaptations that enable therapeutic tolerance, including anti-apoptosis signaling, DNA damage repair, and alterations to cell cycle progression.

Cells may evade drug action through structural alterations of the target [13,14] and up– or downregulation of its expression [24]. Similarly, pathway bypass can be achieved either by reactivating the target pathway through downstream alterations or by activating compensatory pathways that bypass the targeted pathway entirely [25,26]. Drug efflux through ATP-binding cassette family transporters can confer a multidrug-resistant (MDR) phenotype and has been observed across cancer types [27,28]. Tumor cells may also destroy or inactivate drugs or prevent the conversion of a prodrug to its active form [29]. Further, quiescence is a necessarily costly resistance mechanism to cell-cycle-dependent therapeutic pressures; by halting progression, these cells incur a loss of proliferative capacity but establish dormancy-like maintenance programs to withstand treatment or evade the immune system [30]. The persistence of these dormant cells through treatment holiday or remission can contribute to relapse and are a notable departure from the models discussed here that simplify the relationship between cell proliferation and population viability [31].

At the population level, treatment failure and recurrence can also result from inadequate drug delivery due to structural or spatial variation across tumors [32,33]. Intratumoral heterogeneity, driven by genomic instability or divergent fitness landscapes, may further drive diversification and lead to the generation of resistant clones as described above [34]. Because cancer cells can employ multiple resistance mechanisms either concurrently or sequentially [35], durable therapeutic control of tumors remains difficult to achieve. Thus, treatment strategies that explicitly incorporate eco-evolutionary principles for population management could hold promise in preventing resistance emergence.

### Utility of Evolutionary Strategies for Cancer Management

The eco-evolutionary framework of cancer was first conceptualized by Cairns and Nowell in the mid-1970s [36,37]. Concurrently, other fields were actively addressing the consequences of selection under therapy. Resistance management strategies were developed to prevent the spread of pesticide resistance in the early 1970s [27] and later adapted for infectious diseases through antibiotic stewardship initiatives in the 1990s [38]. These guidelines have since been proposed as universal principles of therapeutic resistance management with potential utility in cancer therapy [8,39], as we recognize natural selection as fundamental to cancer progression [18,40].

Adaptive therapy (AT), a resistance management strategy for cancer, was proposed by Gatenby and colleagues in 2009 [4]. Unlike fixed-dose regimens, AT dynamically adjusts treatment intensity based on measured tumor burden, aiming to maintain long-term tumor control rather than maximal tumor eradication [4]. This model assumes two populations: a therapy-responsive, sensitive cell population and a therapy-resistant cell population. It also assumes that there is a cost to therapeutic resistance, such that in the absence of therapy, sensitive cells are expected to out-compete resistant clones. AT leverages therapeutic intervention as negative pressure against sensitive clones, and competition from sensitive clones as negative pressure against resistant clones, enabling long-term therapeutic management of cancers [41–43,4].

## Explanatory Box 2. Theory of Adaptive Therapy

Adaptive therapy leverages competitive interactions between therapy-sensitive and therapy-resistant clones to keep populations of resistant clones at a controlled, low level. As previously stated, MTD approaches exert negative selection on drug-sensitive cells; in the absence of competitive pressure, resistant clones are free to proliferate and drive clonal expansion. This phenomenon is termed competitive release [44].

Adaptive therapy largely relies on the assumption that maintaining a resistant phenotype confers an intrinsic fitness cost to the cell, and that resistance is adaptive only under therapeutic pressure. In the absence of pressure, resistant cells are outcompeted by fitter sensitive cells. By modulating dosage to tumor burden (estimated by imaging or biomarkers), AT forces tumor population dynamics into predictable, controllable cycles (Figure 2). In recent years, preclinical and clinical data have demonstrated the success of AT in increasing time to progression, improving overall survival, and reducing drug toxicity [41–43,45,46].

**Figure 3.**
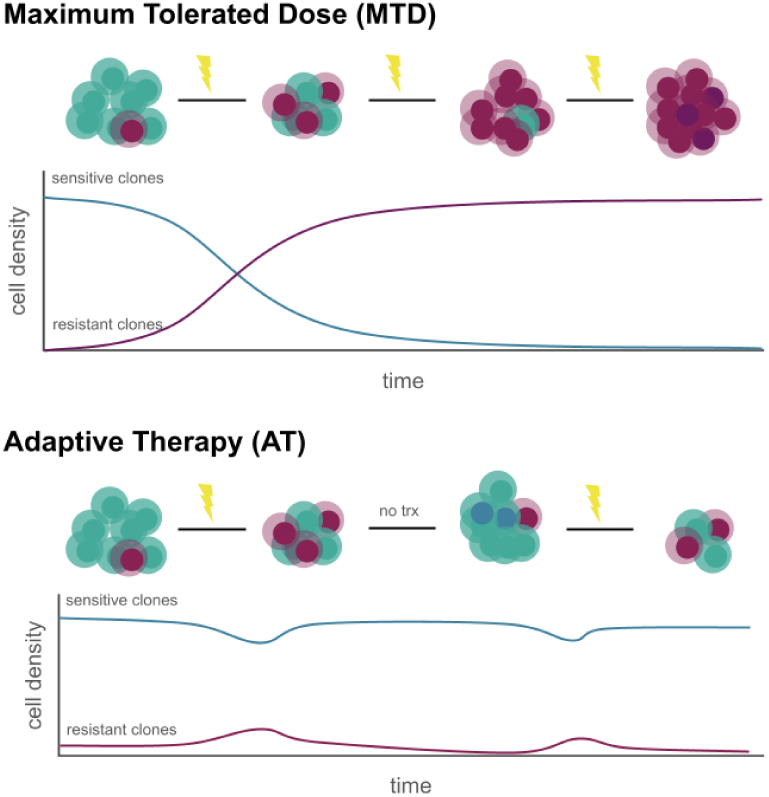
Adaptive therapy vs. Maximum Tolerated Dose. **Fig. 3** Constant dosing that does not modulate in accordance with tumor response may select for resistant cells. Adaptive doses scale with tumor response to maintain a population of therapy-sensitive cells to suppress the resistant population. Cycling adaptive doses can enable long-term tumor control. Adapted from Zhang et al. (2017).

Valcz et al. suggest that AT strategies may be impeded by non-genetic resistance mechanisms [47]. Such mechanisms may enable the rapid expansion of a resistant clonal population at minimal cost. However, non-genetic resistance mechanisms may still be costly and stable enough to be selected upon [10,11,20]. The point stands, however, that dynamic phenotypic switching can quickly deplete the reserve of sensitive cells, which are leveraged to stifle the growth of resistant clonal populations. Conversely, mathematical models have proposed that AT need not rely on a cost of resistance to be successful, and that regular cell turnover can increase the time to progression in AT relative to MTD in tumors approaching carrying capacity [48].

Clinical adoption of AT remains limited. This is largely due to the need for stronger clinical evidence and optimized treatment protocols. Early clinical trials have demonstrated the potential of AT in managing reproductive cancers and highlighted challenges in neuroendocrine-type tumors [43,49,50]. Further research is necessary to identify tumor genotypes and phenotypes most amenable to AT and to refine patient-specific treatment parameters in the clinical setting [51].

Intermittent strategies also rely on the presumed cost of resistance. For instance, intermittent androgen deprivation therapy is commonly administered at periodic doses to reduce comorbidities and delay androgen insensitivity in prostate cancer [52]. The underlying belief is that androgen insensitivity is so costly to individual cells that this characteristic will not emerge *en masse* without significant selective pressure; this idea has some credence but is not well-established in clinical data [53]. Similarly, metronomic therapy can improve tumor control relative to continuous MTD administration [54]. Some therapy-resistant tumors may regain sensitivity following a drug holiday, consistent with the idea that resistance is selected against in the absence of therapy [55]. However, it follows that resistance characteristics with little to no fitness cost are likely to persist even in the absence of treatment, as negative selection against these clones is weak [56]. It is therefore pertinent to understand the predictors of costliness in resistant tumor cells to better stratify patient response to a given therapy and predict the longevity of treatment efficacy.

### Generality of the Assumption of Resistance Cost

Life history theory predicts that investment in defense requires reallocation of resources away from growth and proliferation [57,58]. A model by Hausser et al. proposes that cancer cells divert resources among five distinct “universal cancer tasks”, with investment constrained by tradeoffs along a continuum of strategies [59].

Further, resistant clones are often present in the treatment-naive tumor at diagnosis, but typically at frequencies so low as to be rarely detected [15,60,61], suggesting that these lineages do not have a fitness advantage over sensitive clones. Cell population sizes are so large, and mutation rates are so high in cancers [40], that they are likely to spontaneously generate resistance mutations, perhaps many times within the same cancer. This perspective suggests that some cases of primary resistance may involve a resistant clone with a fitness advantage that expanded to fixation before diagnosis, whereas acquired resistance implies a fitness cost.

Across biological systems, resistance management strategies have proven successful because the functional compromises they impose render resistance costly. Fitness costs of resistance have been documented in insects [62], plants [63,64], viruses [65,66], fungi [67], bacteria [56], and parasites [68,69]. In oncology, the assumption of resistance cost is frequently embedded in mathematical models of tumor dynamics [4]. Experimental successes of adaptive therapy, intermittent dosing, and drug-holiday strategies are often attributed, at least in part, to these costs [4,43,45,70]. Given pervasive biological constraints, the existence of resistance-associated tradeoffs is a reasonable and widely invoked assumption.

However, to date, no comprehensive systematic review has evaluated whether therapeutic resistance generally entails a fitness cost in the absence of therapy across drugs, cancer types, etc. Currently, there is limited consensus regarding which resistance mechanisms are likely to be costly, the conditions under which these costs emerge, or how environmental context modulates their magnitude. Such costs can be evaluated using *in vitro* and *in vivo* competition models between sensitive and resistant cell populations. We aim for this research to support drug development and refine cancer management strategies. [45,51].

## Methods

### Search Strategy

This systematic review was conducted in accordance with established guidelines, including PRISMA2020, to define retrieval, screening, and data collection protocols [71,72].

Given the narrow scope of the research question and the variability of keywords used in the literature, a set of thirteen publications identified through a non-systematic search was used as a positive control to inform the construction of itemized search queries. Searches were performed in the databases Scopus, PubMed, and Google Scholar, with database-specific adaptations made to accommodate query language constraints. All search strings are provided in Supplemental Table S1.

Final database searches were conducted on 3 October 2025, yielding 85 records from Scopus, 58 from Google Scholar, and 37 from PubMed. Articles that do not contain original research were filtered out. To supplement database retrieval, the large language model-assisted literature database SciSpace was used to identify additional relevant publications, returning 50 peer-reviewed articles from its associated library. The SciSpace search query is listed in Supplemental Table S1. Citation searching was employed but yielded no additional eligible studies.

### Inclusion Criteria and Screening

After initial retrieval, duplicate records, abstracts or titles that did not meet relevance criteria (therapeutic resistance in cancer), publications that were not primary research reports, and records that could not be retrieved were excluded (Supplemental Figure S1). The remaining 122 papers were screened according to the following eligibility criteria: (i) experiments depict direct competition (two distinct cell lines in direct contact in a mixed culture), (ii) between therapy-resistant and therapy-sensitive cancer cell lines, (iii) in the absence of therapeutic pressure. These criteria are designed to capture experimental outcomes of direct interaction between cell lineages with differential responses to a particular therapy. Among relevant studies, 23 were excluded for failing to meet all three experimental criteria.

After preliminary full-text eligibility screening, the 45 preliminary papers considered for inclusion were processed through SciSpace with a plain-language prompt (Supplemental Table S1) to search for publications not already captured. No additional papers were retrieved through this step. Final screening measures were conducted by evaluating the experimental design and reviewing supplemental figures. This process led to the exclusion of 22 additional publications. Ultimately, 23 publications that satisfied all inclusion criteria were advanced to the data collection and analysis step.

### Data Collection and Analysis

For each eligible experiment, twenty-eight descriptive features were extracted and recorded (Supplemental Table S2). Certain features relevant to our analysis require inference due to variation in study design. We define fitness using population growth rate as a proxy for reproductive competency and survival [73], and subsequently define the winning lineage as that having the greater population growth rate in coculture at experiment endpoint. Further, we define the resistant lineage as one that tolerates higher exposure to therapy than its sensitive counterpart.

Data items that could not be identified in the main text or within a direct citation were recorded as not available (NA). The complete table of included publications is provided in Supplemental Table S3. Statistical analysis of the association between features and the cost of resistance was performed using Fisher’s exact test, implemented via the stats package in R (version 4.5.1).

## Results

### Resistance Characteristics Predict Costliness

Of 49 competition experiments sourced from 23 peer-reviewed publications that met the inclusion criteria, resistant cells won in 13 (26.5%), lost in 33 (67.3%), and had neither an advantage nor a disadvantage in 3 (6.1%). For each experiment, we recorded data on experimental design and model features. The majority of studies investigated resistance to chemotherapeutic agents (n=24), though targeted therapies remain the second largest category (n=18). A small number included resistance to immunomodulatory drugs (n=7) or gamma radiation (n=2). We did not identify any studies employing hormone therapy-resistant lineages. Cancer types included breast, ovarian, cervical, prostate, colorectal, liver, lung, and hematopoietic cancers. A wide variety of methods were used to generate resistant lineages, though in all but 4 cases, the resistant population was derived from the parental sensitive population. Excluding these studies from subsequent analyses did not significantly change the results (Supplemental Figure S2).

We assessed ten features for association with experimental outcome: model type, method of generating resistant cell lines, whether the resistant line was derived from the sensitive line (isogeny of populations), cancer type, category of resistance mechanism employed, specific pathway implicated by the resistance mechanism, treatment category, drug class, whether the experiment was resource-limited *in vitro*, and model organism (Figure 4). The only predictor that reached significance threshold was the mechanism of resistance employed by the resistant population (p=0.0154).

**Figure 4.**
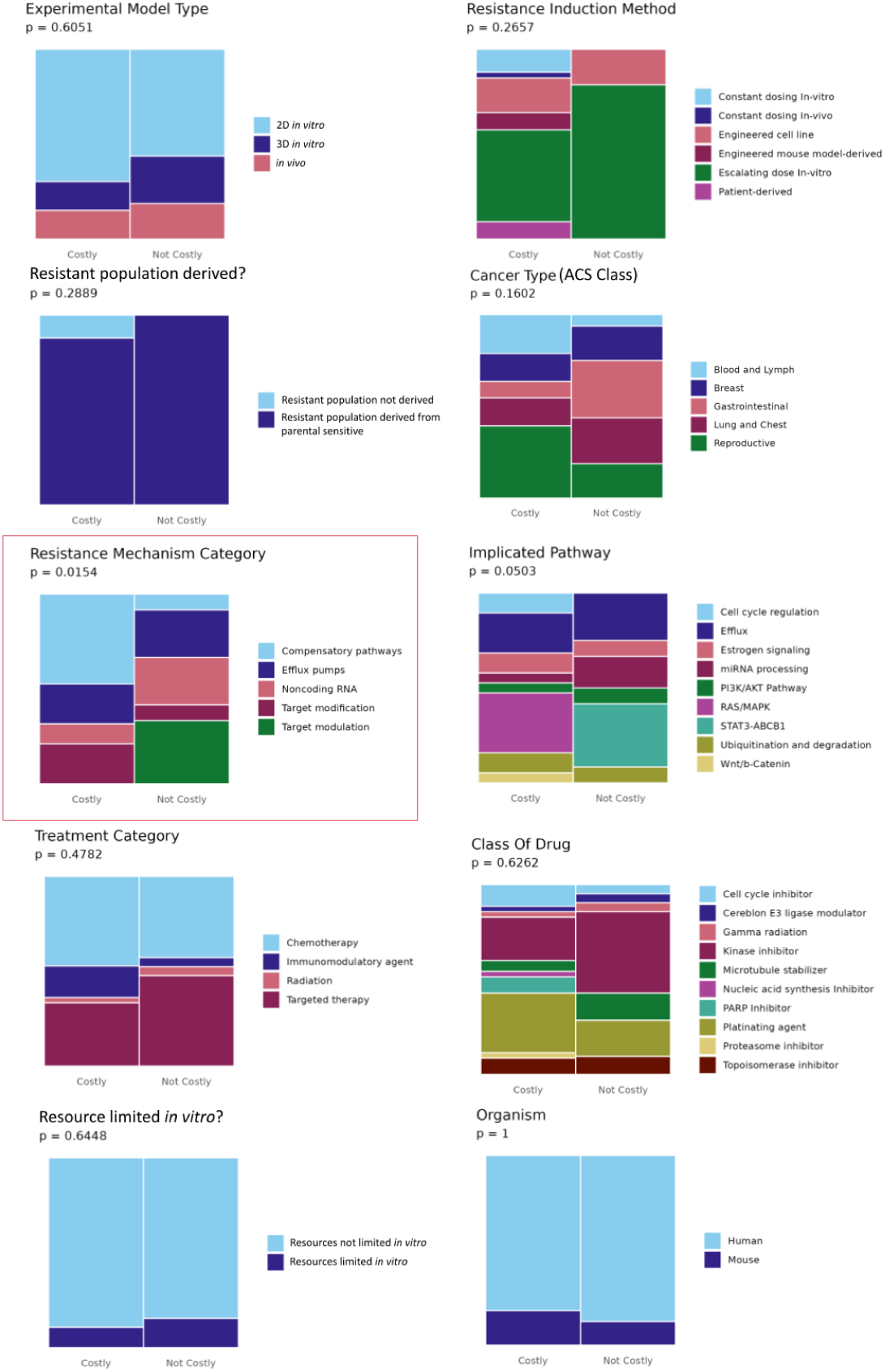
Outcome by Recorded Feature. **Fig. 4** Mosaic plots depict distribution of experiment outcome among 10 experimental features. In order with associated column number from Supplemental Table S2: Experimental model type (Col. 5), resistance induction method (Col. 22), resistant population derived? (Col. 12), cancer type (Col. 11), resistance mechanism category (Col. 18), implicated pathway (Col. 20), treatment category (Col. 23), drug class (Col. 24), resource limited *in vitro?* (Col. 9), organism (Col. 10). Column height indicates proportion among group (Costly n=33, Not Costly n=16). Not Costly includes instances where resistant cells won in competition or no significant fitness gap was observed in competition.

### Target Modification

Three studies in hematopoietic malignancies demonstrated that resistant cells evaded therapy by altering the target protein. Knockout of *CUL4B*, a gene encoding a subunit of E3 ubiquitin ligase, confers robust resistance to lenalidomide in multiple myeloma. *CUL4B* KO clones had a growth penalty in coculture against parental cells, suggesting that disruption of the ubiquitination pathway incurs a cost for resistance that is not adaptive in the absence of drug [74]. However, the same study showed that mutational alteration of the target transcription factor *IKZF1* does not confer a fitness disadvantage to resistant cells in competition [74].

Further, resistance to the targeted proteasome inhibitor bortezomib in multiple myeloma is conferred by a missense mutation in *PSMB5* near the drug-binding pocket. Alterations in the active site of this proteasome subunit may reduce catalytic efficiency, but proliferation was not significantly affected [75]. Though this mechanism of bortezomib resistance appears to confer little autonomous fitness penalty, PSMB5 A20T mutants were outcompeted by sensitive clones in untreated coculture [76,77]. These data suggest that bortezomib resistance via proteasome-binding-site modification may incur a fitness penalty that sensitive cells exploit.

Target modification may prove costly in breast cancer as well. Resistant cells can escape treatment by the cell-cycle inhibitor NU6102 via mutational modification of cyclin-dependent kinase 2 (CDK2), the target of NU6102. Subsequently, a lesser proportion of resistant cells was observed in S phase at any given time, indicating cell-cycle alterations between clones and supporting the overall lower competitive fitness of the resistant line [78].

### Non-coding RNA

Long non-coding RNAs (lncRNA) and microRNAs (miRNA) mediate cell proliferation, motility, and physiology; dysregulation of these systems has been investigated as initiators and drivers of cancer progression [79]. Radiotherapy resistance in PC3 prostate cancer cells was driven by miRNA-95, which drove the activity of sphingolipid phosphatase [80]. These radiotherapy-resistant clones bore a higher proliferation rate than sensitive clones in both monoculture and dominated spheroid coculture [81]. Success in competition and aggressive phenotype suggest that this mechanism of resistance does not confer a significant fitness cost. Conversely, in models of the prostate cancer line DU145, the radiotherapy-resistant population was driven down by the presence of sensitive cells despite an intrinsic growth advantage. High abundance of the lncRNA UCA-1 was noted in resistant DU145 clones, a mediator of growth and proliferation [82]. Unlike miRNA-95-overexpressing clones, however, the UCA-1-high clones had a fitness disadvantage in competition. This disagreement highlights the differential costs of various resistance mechanisms.

Further, cross-spectrum resistance in non-small cell lung cancer (NSCLC) was conferred by stochastic mutations in *Dicer1*, a key gene involved in miRNA processing. Multidrug-resistant *Dicer1* mutants (“M1”-”M3”) demonstrated comparable growth rates to parental clones when cultured separately. The resistant “M1” mutant was outcompeted by sensitive clones across a range of seeding densities, but the “M2” and “M3” mutants dominated in coculture [83].

### Target Up– and Downregulation

RPTK overexpression appears to confer a growth advantage and resistance to the multikinase inhibitor lenvatinib in HCC, even in the absence of therapy. Accompanying this is a marked shift towards glycolytic metabolism. Resistant clones dominate both *in vitro* competition assays and mixed tumors in mouse models. However, tumors composed solely of resistant cells were smaller in volume than those composed solely of sensitive cells, suggesting a potential hurdle to the altered metabolic program in a physiological environment [84].

### Efflux Pumps

In breast cancer, clones resistant to the topoisomerase inhibitor doxorubicin displayed an efflux pump-high, glycolytic phenotype. However, the expense of maintaining this resistant phenotype varied; it proved either disadvantageous or beneficial in coculture between studies, despite increased resource demand and variable intrinsic fitness consequences [34,85]. In a hepatocellular carcinoma model, efflux pump overexpression conferred resistance to oxaliplatin and a competitive benefit in two-dimensional coculture [86].

Further, the PARP inhibitor olaparib disrupts DNA repair and is particularly effective in BRCA-mutant cancers [87]; however, resistance often arises through efflux pump overexpression [88,89]. Olaparib-resistant ovarian cancer lines exhibited epithelial-mesenchymal transition (EMT) markers and upregulation of efflux pumps. In BRCA1-proficient cell lines, sensitive and resistant clones had similar growth rates when cultured separately; in BRCA1-deficient cell lines, however, the olaparib-resistant clones had an intrinsic fitness penalty [90]. In both cell lines, resistant cells were considerably outcompeted by sensitive cells [27,90]. Taken together, these studies demonstrate that the intrinsic costliness of maintaining an efflux pump-high MDR phenotype in ovarian cancer may confer a fitness disadvantage in competition, particularly when it is associated with an EMT phenotype.

### Compensatory Pathway Activation

A panel of *NRAS* mutations renders myeloproliferative neoplasms resistant to ruxolitinib, a JAK inhibitor. Mutations in *RAS* are among the most recognized oncogenic drivers across cancers and offer cells an opportunity to recover fitness deficits induced by JAK inhibition by employing non-JAK/STAT-mediated signaling pathways instead [91]. Despite pathway bypass conferring resistance, ruxolitinib-resistant cells were outcompeted by sensitive cells in drug-free conditions both *in vitro* and *in vivo*. The same was seen in EGFR-inhibitor-treated lung cancer models, where compensatory reliance on oncogenic KRAS conferred a competitive disadvantage [92]. Similarly, mutations in the *Apc* tumor suppressor gene unconstrain *Wnt* signaling, another oncogenic driver across cancers [93,94]. Cells dependent on accelerated *Wnt* signaling are negatively selected in the absence of therapy and are outcompeted by sensitive cells *in vivo* [95]. This phenomenon is termed oncogene overdose: cell death or growth arrest triggered by overreliance on an aberrant oncogenic signaling pathway [96]. In these studies probing *RAS* and *Wnt* mutations, oncogene overdose may contribute to decreased fitness in the absence of therapy [97].

Resistant breast cancer cells compensate for the cytotoxicity of ribociclib, a CDK4/6 inhibitor, by increasing estradiol production and driving growth signaling. Both sensitive and estradiol-overproducing resistant clones have comparable growth rates when cultured separately. However, resistant cells in coculture promote sensitive cell growth through secreted estradiol production, resulting in a net fitness advantage to the sensitive population and subsequent suppression of resistant cells [98].

Comprehensive review of the fitness consequences of specific resistance characteristics is limited by the small number of studies present in each category, as well as the lack of an explicit resistance mechanism described in eight of twenty-three studies. However, general principles appear to underlie competition between therapy-sensitive and therapy-resistant clones reliant upon similar resistance characteristics across cancer types and treatment pressures. This significant association between mechanism of resistance and relative fitness has been demonstrated previously in antimicrobial resistance literature [56], and we are excited to report trends that may suggest the same in cancer.

### Other Trends

Though not meeting the significance threshold, the association between experiment outcome and implicated pathway approaches significance (p=0.0503). In particular, all observations of RAS/MAPK pathway alterations contributing to the resistant phenotype bore reduced competitive fitness (n=6). Conversely, alterations to STAT3-dependent transcriptional pathways tended not to incur costs (n=4). Similar to the mechanism of resistance, this association is somewhat intuitive, providing mechanistic insight into the changes observed in resistant lineages. This pattern may be encouraging in identifying the tradeoffs incurred by specific pathway alterations.

In our review, the experiment modality did not significantly influence the likelihood that resistance had a fitness cost (p=0.6051). However, population dynamics may shift between models. In one case, a fitness advantage observed *in vitro* for the sensitive population was absent in the corresponding *in vivo* trial [99]. Further, three-dimensional models illustrate spatial and colocalization dynamics, which can modulate hypoxia, extracellular architecture, and nutrient availability [100]. Bacevic et al. demonstrated AT efficacy in a three-dimensional colorectal spheroid model, but not in two-dimensional culture [78]. Paczkowski et al. found that in cocultured prostate spheroids, highly proliferative resistant cells constituted much of the periphery, while sensitive parental cells were confined to the hypoxic spheroid core [81]. Similarly, Thege et al. found that drug flux to the sensitive spheroid core was reduced by surface localization of resistant clones [101]. Spatial dynamics are therefore critical in assessing fitness landscapes relevant to clonal competition. Some studies present *in vitro* experiments designed with this limitation in mind, and demonstrate that interactions with stromal cells and competition for limited space, oxygen, and nutrients influence the competitive ability of clones [41,81,85,102].

The lack of a cost of resistance in a third of cases does not appear to be explainable by the abundance of resources in the coculture experiments. Two of the five experiments that restricted resources *in vitro* still found no evidence of a cost of resistance, though in all of the remaining experiments, they did find that resistant cells that could win competition in rich media lost competition in resource-restricted media [41].

Also surprising to us was the weak association between the method of generating the resistant population and experimental outcome (p=0.2657). A seminal study by Rath et al. found that altering treatment regimens produces resistant clones with distinct mechanisms of resistance and doubling times that varied nearly sevenfold between clones. Replicates exposed to stepwise increases in drug concentration developed resistance via gene amplification of the dihydrofolate reductase gene and had faster doubling times than those derived from constant exposure to lethal drug concentrations, which lacked gene amplification [103]. Subsequent work has established that the mutational background and expression profile of the parental line, the dosing regimen, and the drug-free maintenance period influence the resistance characteristics of the selected clones [104]. Given the possibility that populations can develop distinct resistance mechanisms in response to the drug microenvironment, we do not preclude that this may influence cell fitness, despite the weak association identified here.

We were interested in evaluating whether experimenters quantified resistance and, if so, if the magnitude of resistance was predictive of competition outcome. It may reasonably follow that a lineage expressing a high degree of resistance to therapy (say, twenty-fold compared to the sensitive population) would incur different fitness compromises than one that bore two-fold resistance compared to the sensitive population. This may simply reflect divergent phenotypic optima under different environmental conditions [105]. Performing a log-regression analysis using experimental outcome (Costly/Not Costly) and the quantitative fold-change in IC50 values between sensitive and resistant cell lines, we did not find a significant association (p=0.535, Chi-Square Test) (Supplemental Figure S3). Just over half (n=26) of the experiments quantified resistance by comparing IC50 values across lineages, limiting the analysis’s statistical power.

### Autonomous and Competition-dependent Fitness Consequences

Fitness measures in monoculture often differed from fitness measures in coculture (Figure 5). Significant gaps were observed more often in coculture than in monoculture, where 14 models did not identify a significant difference in growth rates between populations; all of these studies demonstrated fitness gaps between populations in coculture. The data support the reasoning that autonomous fitness differences are exacerbated due to competitive interactions. Additionally, there were a small number of cases where fitness gaps were reversed in competition. Resistant cell lines may grow at higher rates than sensitive cell lines when cultured independently, but suffer fitness deficits in competition [81]. Conversely, the opposite – that a slower-growing resistant population gains an additional growth advantage from coculture – was true in some cases as well [84,85]. 18 of 49 experiments did not supply monoculture growth data.

**Figure 5.**
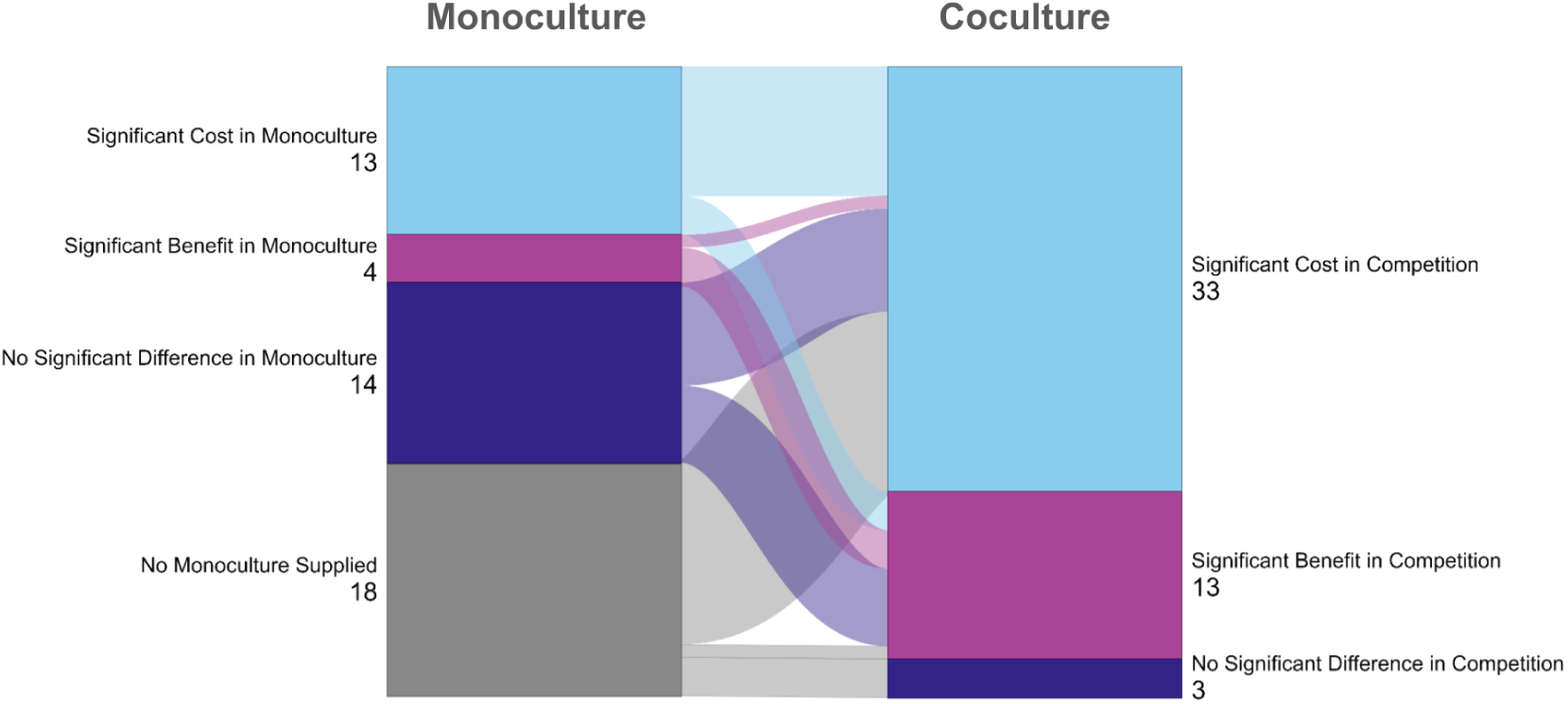
Outcomes of Competitive Cocultures and Homotypic Monocultures. **Fig. 5** Experiment outcomes differed between mono– and coculture in 18 of 31 cases. 10 instances where resistance proved costly in monoculture also indicated a significant cost in coculture. Conversely, growth benefits for the resistant line were observed both in mono– and coculture in just 3 studies. All 14 experiments that demonstrated no significant growth rate discrepancies between populations demonstrated significant gaps in competition; 6 showed that resistant populations had a growth advantage in competition, while 8 showed that resistant populations incurred a fitness cost in competition.

Measuring population fitness in monoculture is critical to understanding nuanced competitive outcomes. Cell-autonomous fitness consequences impose a burden on the cell regardless of the presence of competitors in the microenvironment. On the other hand, competition-dependent, or non-cell-autonomous, fitness consequences are modulated by interactions between cells; the autonomous fitness gap between cell types and relative population densities are relevant features, and game-theory models can describe their interactions [102,106]. The competitive Lotka-Volterra model, an ecological model of competition, considers both autonomous growth rates and carrying capacities for each population as well as their competitive effects on each other [107].

Data from monoculture models can supply growth rate and carrying capacity for the sensitive and resistant cells. This informs the use of ecological frameworks to describe and model interactions between populations in competition. Divergence from the predictions of the Lotka-Volterra model can reveal complex competition-dependent fitness consequences in coculture, which include facilitation, frequency-dependent selection, phenotypic shifts, and cell-cell fusion. These interactions have the potential to modulate cost of resistance [92].

**I. Facilitation**. The ecological definition of facilitation broadly includes all interactions that result in a fitness benefit for at least one participant [108]. Antagonism describes interactions where one population benefits at the explicit expense of another. Under drug-free conditions, sensitive cells can derive an additional fitness benefit from the presence of “loser” resistant lines, mediated by cell-cell contact or paracrine signaling [109,110]. Conversely, resistant clones can derive an additional fitness benefit from competition, partially characterized by upregulated glycolytic metabolism absent in populations grown in monoculture [84].
**II. Phenotypic shifts in drug response**. Phenotypic shifts modulate the fitness differential between clones and drive the population toward homogeneity under drug pressure in two major ways: by improving the response of resistant cells to therapy (resensitization) or increasing the tolerance of sensitive cells to therapy (desensitization). Desensitization of sensitive clones by the presence of resistant clones was observed across various competition models. The ability to transiently adopt resistant phenotypes has significant implications for the success of adaptive therapy [47]. However, the cost assumptions for these phenotypic transitions differ mechanistically across models; for instance, microvesicle-mediated horizontal transfer of drug efflux pumps confers a fitness benefit to treatment-naive cells under therapeutic pressure [111]. However, the cost of maintaining the resultant MDR phenotype is borne by the recipient due to the significant metabolic burden imposed by efflux pumps. Sensitive cells in this model return to baseline drug tolerance levels upon treatment cessation, shedding the high-cost phenotype. Alternatively, tolerance can be induced in the sensitive population by paracrine signaling of growth factors from resistant cells. In this model, sensitive cells benefit from growth factor production in the microenvironment without incurring a cost, while the donor population bears the cost [98,112]. Craig et al. also propose a phenomenon termed cooperative adaptation to therapy (CAT), wherein resistant clones appear to induce phenotypic transitions in sensitive lineages, exceeding the tolerance observed in monoculture, through direct cell-cell contact [83]. The proportion of resistant cells in the total population has been suggested to significantly modulate population drug response [113]. Alternatively, resensitization of clones was also observed in coculture. Secreted components from sensitive clones can reduce the competence of resistant clones under therapy [86,101]. Interestingly, one model also demonstrated that sensitive clones may maintain their fitness advantage even under sublethal therapeutic conditions [86]. Taken together, this may suggest that a significant density of therapy-sensitive clones may both exert antagonism against resistant clones and induce phenotypic transitions that favor a sensitive state. This provides further rationale in some cases for the maintenance of a population of sensitive clones to combat drug resistance acquisition.
**III. Fusion**. Cell-cell fusion is similar to, but mechanistically distinct from, phenotypic transition, as it is irreversible and results in the effective removal of constituent cells from the population. Cell-cell fusion is regarded as a potential hallmark of cancer progression initiated by microenvironmental pressures [114]. In an NSCLC model, novel cell-cell nuclear fusion occurred at high rates under drug selection between sensitive and resistant clones, despite a steep competitive edge against chemoresistant clones in the untreated environment. The fused cell exhibited greater chemoresistance across a spectrum of agents than the parental chemoresistant clone, suggesting not only a novel adaptation to therapy but also an acceleration in the evolution of the mixed population due to genetic heterogeneity [115]. The propagation of drug-resistant clones, however, is stifled by the presence of sensitive clones in the absence of therapy; it still stands to reason that even in these rare fusion events, conservative application of therapy could maintain a significant pressure against resistant clones that may drive the resistant population down.
**IV. Density-Dependent Selection.** Clonal populations exert competitive pressure according to their relative densities. In one model, alectinib-resistant and sensitive NSCLC parental clones have similar growth rates when cultured separately. In coculture, however, competition is modulated by drug presence, cancer-associated fibroblasts, and population density. In all conditions but a portion of those cultured with fibroblasts and no drug, parental growth rate is outpaced by that of resistant clones [84]. The major shifts in the growth rates of sensitive and resistant clones observed reinforce that cell-cell interactions can influence the success of phenotypically distinct clones in competition. We identified one case where the winning clone was that which was present in the greater proportion from seeding [41]. It has been suggested that frequency dependence can ameliorate the cost of resistance and may be in part to blame for rescue subpopulations that contribute to relapse, or for pre-existing resistant clones [92].

## Discussion

Fitness of a mutation or phenotype is a complex phenomenon, dependent on the environmental context, the genomic background, and the competitors (as well as the cooperators) in that environment. The experimental tests of the fitness of resistant clones to date are too few to control for variation introduced by different genomic backgrounds, different tissues of origin, and varying microenvironments. By grouping studies by the different mechanisms of resistance, we identified fitness patterns across the 49 competition experiments we analyzed. In addition, the experiments highlighted several important complexities in measuring the cost of therapeutic resistance.

Our primary finding that the mechanism of resistance is most closely associated with competitive fitness is perhaps unsurprising, mirroring patterns in antimicrobial resistance [56]. Antineoplastic therapies target essential cell functions. To evade death by therapy, those essential functions must be compromised in some way; therefore, alterations, whether mutational in nature or not, are likely to impose a fitness consequence. We found substantial variation between drugs and cell lines; however, some combinations showed that resistance conferred a competitive advantage even in the untreated environment. We noted similar resistance adaptations emerging under different drug pressure (e.g., MDR via efflux pumps) as well as dissimilar costs arising from resistance to the same therapeutic agent. (e.g., distinct resistance mechanisms in radioresistant prostate cancer, one costly in competition, and one not [81]). Further work is needed to understand the proclivity to particular evolutionary fates under drug pressure, underscored by our weak association between treatment category and cost (p=0.4782).

The results presented here are most vulnerable to challenges to the notion of the cost of resistance itself. Lenormand et al. present a thorough exploration of the concept of costs and primarily raise issues regarding the relationship between the non-ancestral environment and population fitness within it. Lenormand rightfully objects to the notion of a single cost; there are as many costs as there are environments (some of which are not negative fitness consequences at all). Further, the population may never reach the phenotypic optimum in the ancestral environment and instead be better acclimated to a novel environment [105]. Thus, assessment of fitness in a single drug-free environment may fail to capture the full spectrum of pressure exerted by various drug concentrations, under which the optimum phenotype may vary significantly.

It is difficult to compare results across experimental systems and even across drugs. Though we did not find an association between the degree of resistance and whether or not resistance appeared costly, it is not clear that a two-fold change in the concentration of a cytotoxin that kills 50% of the cells is evolutionarily equivalent to a two-fold change in the concentration of a targeted therapy that kills 50% of the cells. The metabolic costs of those forms of resistance may differ significantly, which is why we also considered an interaction between the degree of resistance and its mechanism.

This therefore calls into question the viability of coculture models. If cancer cells are in a culture environment far removed from their physiological microenvironment, and fitness is context-dependent, how can these findings be extrapolated to tumors? It is heartening to know that many of these experiments are indeed embedded in studies evaluating the efficacy of AT or drug-holiday strategies, which, in all cases presented here, corroborate the finding of fitness deficits in coculture [34,41,45,85,99]. While not a one-size-fits-all approach, coculture models capture facets of the tumor microenvironment, the relevance of which can be enhanced by various design choices (e.g., using spheroid models to capture spatial heterogeneity, allowing nutrient and oxygen availability to be nonuniform factors in the microenvironment).

Melnyk et al. highlight cases of antibiotic resistance where the resistant strain persists in the untreated environment despite significant fitness costs, suggesting the following reasons for this observation: (i) linkage between selected loci and resistance loci, (ii) second-line alterations that increase fitness but do not reduce resistance, and (iii) variable pleiotropic effects of resistance mutations [56]. In the selected studies, we identify several additional resistance characteristics that do not constitute proximal resistance mechanisms but nonetheless contribute to cell fitness and competitive ability. Among these are measurable metabolic shifts, EMT markers, and ECM remodeling [41,45,84,86,90]. It is unclear if the proximal resistance mechanism and these secondary resistance characteristics are extricable, but further suggest that widespread phenotypic changes result under drug selection for any of the reasons aforementioned. Though the pleiotropic effects of mutations contribute to fitness and their negative effects may be ameliorated over time, they are distinct from the suboptimal functioning of an altered trait in the untreated environment [105].

### Vulnerabilities and Modulators of Costliness

Fitness is necessarily context-dependent and can vary over space and time [116]. Therefore, it is reasonable to propose that modulation of specific environmental factors might incur greater fitness costs in cells bearing the resistance phenotype, thereby facilitating therapy.

Efflux pumps pose significant drug-delivery challenges across therapy types but also impose a high metabolic burden, which may be leveraged for cross-sensitivity [28,117]. Treating MDR clones with a non-cytotoxic efflux pump substrate (called an ersatzdroge) may increase the fitness cost of the resistant cells [85,118,119]. *In vitro* trials have demonstrated the success of ersatzdroges in exhausting MDR clones [85,118]. However, clinical adoption of ersatzdroge-chemotherapy combination treatments failed to demonstrate significant success in eradicating MDR clones [120,121].

Another concept in evolutionary oncology is evolutionary herding, or the Sucker’s Gambit. This strategy proposes that tumors could be steered into predictable evolutionary states by applying selective pressures that render the population vulnerable to therapy or other exploits [122]. Clinical trials have been proposed to evaluate the efficacy of an initial treatment in enhancing response to a subsequent one, but there have been few tests of this idea [123]. This approach is supported by the possibility that clones at various stages of resistance induction may expose cross-sensitivities that could be exploited by therapy [6,124]. Controlled initiation of oncogene overdose via metronomic therapy has also been demonstrated to prolong tumor control [125].

### Clinical Translation

Our findings corroborate the current understanding of resistance development and present interventional opportunities to delay or halt this process [9,14]. In a recent preprint, Gallagher et al. propose patient-specific “mathematical biomarkers” for predicting AT success, estimating clonal competition from tumor response to an initial cycle of therapy. In conjunction with these sophisticated modeling techniques and measurable biomarkers as a proxy for tumor burden, the mechanistic data we review here can facilitate the development of AT techniques in the clinical setting [126]. The molecular features of resistant cells, particularly if a particular resistance characteristic emerges frequently among treatment-refractory cancers, could be used to monitor the progression of the resistant population.

Therapeutic resistance does not always impose a fitness cost, which presents a challenge for AT; however, a cost of resistance is not the only predictor of success in AT [34,47,48]. It is not clear whether or why AT interventions have different success rates across cancer types, though the microenvironmental context is a promising next step for investigation [127]. We hope that the data presented here may help to explain the between-patient and between-cancer variation seen in the literature.

## Concluding Remarks

Across the 49 competition experiments we identified, a cost of resistance was common but not universal. With a limited sample size, it is challenging to reliably determine the effects of various relevant factors, such as the background genotypes in which resistance evolves, the likelihood of a cost of resistance to any particular drug or therapy, let alone a drug class. A notable disconnect between population fitness in monoculture and coculture demonstrates the need for direct competition experiments to capture clonal dynamics.

In the studies reviewed, we specifically find that employing compensatory pathways may force cells into fitness-compromising states, exposing vulnerabilities that may be exploited. We were surprised to find that structural alteration of the drug target also tended to impose a fitness cost in almost all cases. The fitness consequences of some mechanisms of resistance, such as enzymatic drug inactivation, have not been evaluated and warrant further investigation.

Here, we present empirical evidence on the fitness cost of resistance in cancer and the patterns within our limited purview. We intend for this review to help oncologists address the clinical problem of therapeutic resistance, facilitate the development of therapies that prevent or delay the evolution of resistance, and address open questions in evolutionary oncology [51].

## Supporting information

Table of Reviewed Studies (.tsv)

## Acknowledgements

We would like to thank the Arizona Cancer Evolution Center for supporting this project by providing an avenue for conversation and critique.

## Artificial Intelligence Use Statement

The LLM-assisted platform SciSpace was solely to support literature retrieval and to validate the final list of eligible publications. All outputs were reviewed manually by the authors. No generative artificial intelligence tools, including SciSpace generative AI capabilities, were used in the conceptualization, outlining, drafting, or editing of this publication. All text and figures are the authors’ original and own work.

## Data Availability

The data presented in this review are available in the article and in its online supplementary material.

---

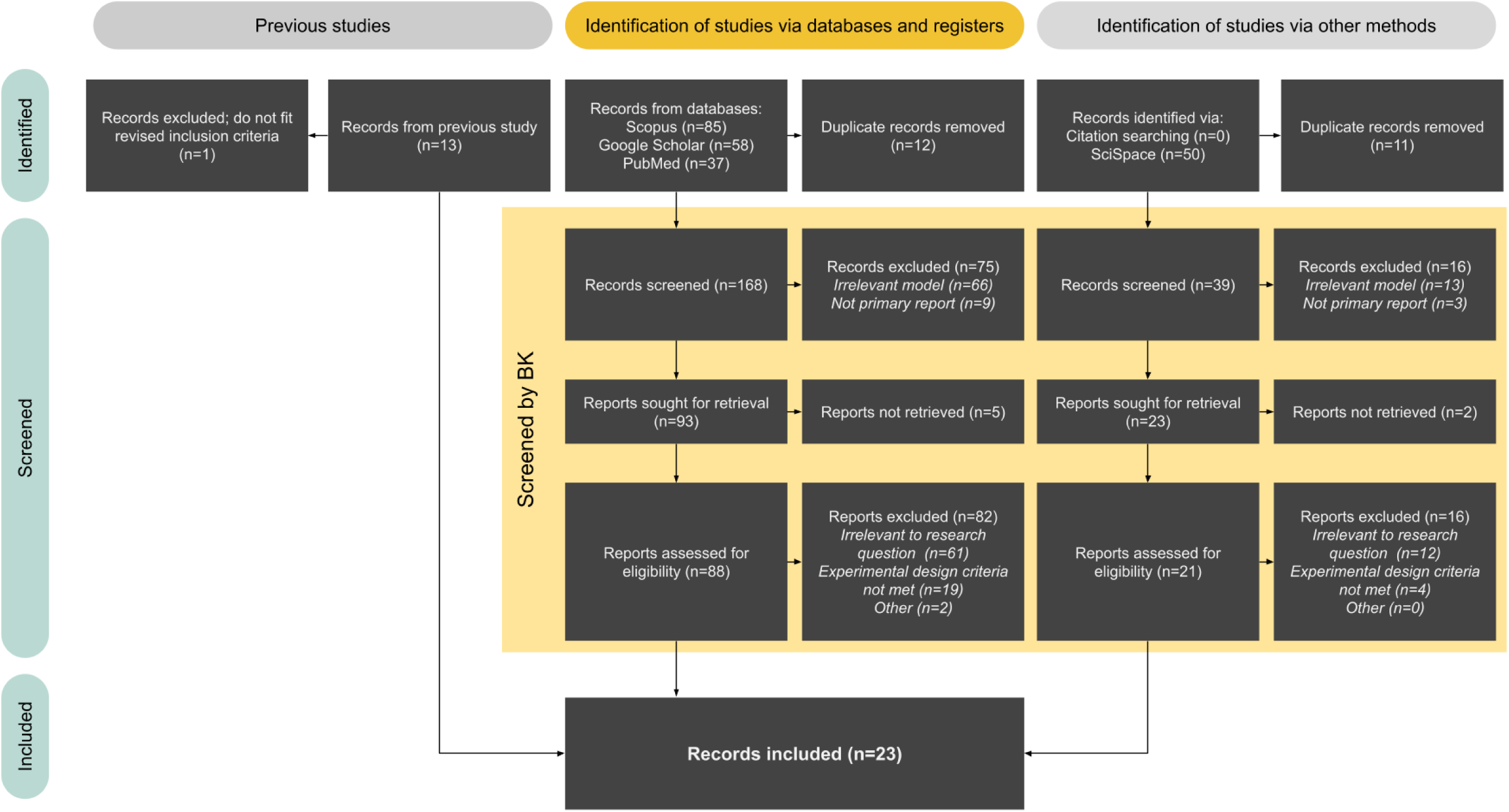
SUPPLEMENTAL FIGURE S1A.

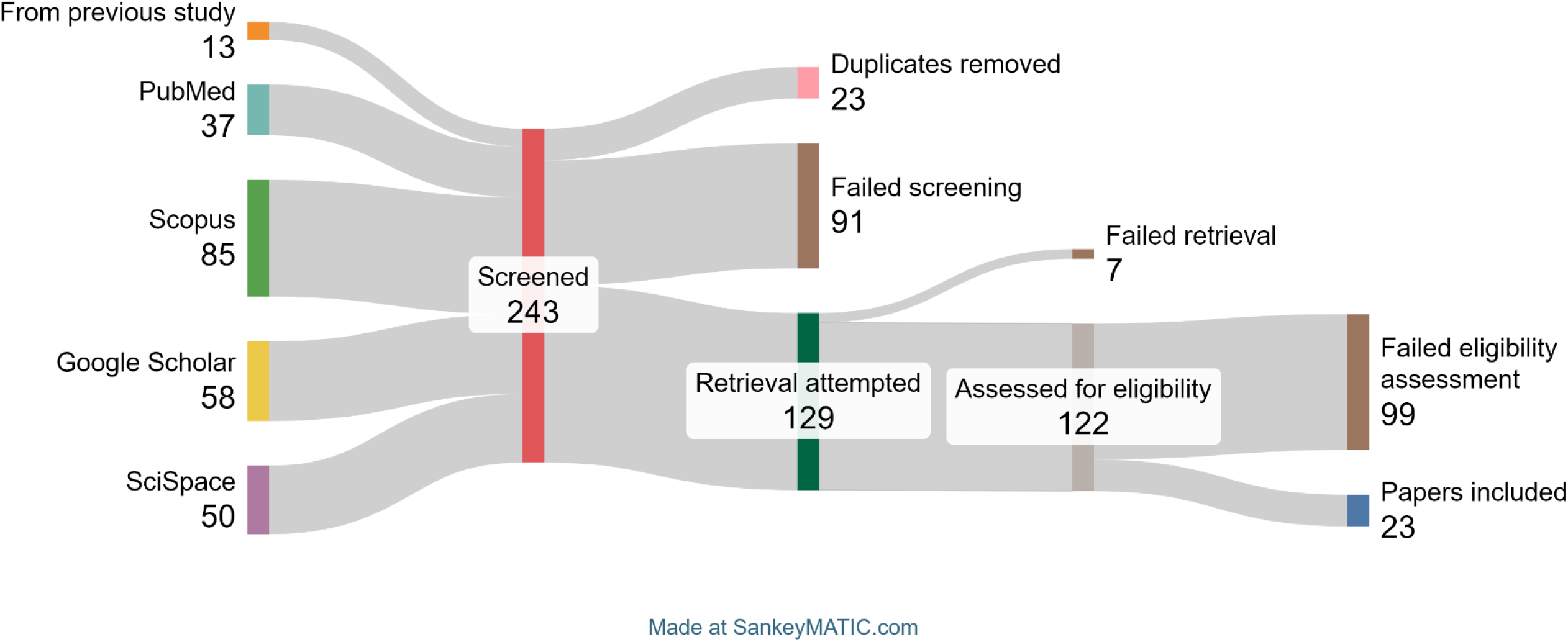
SUPPLEMENTAL FIGURE S1B.

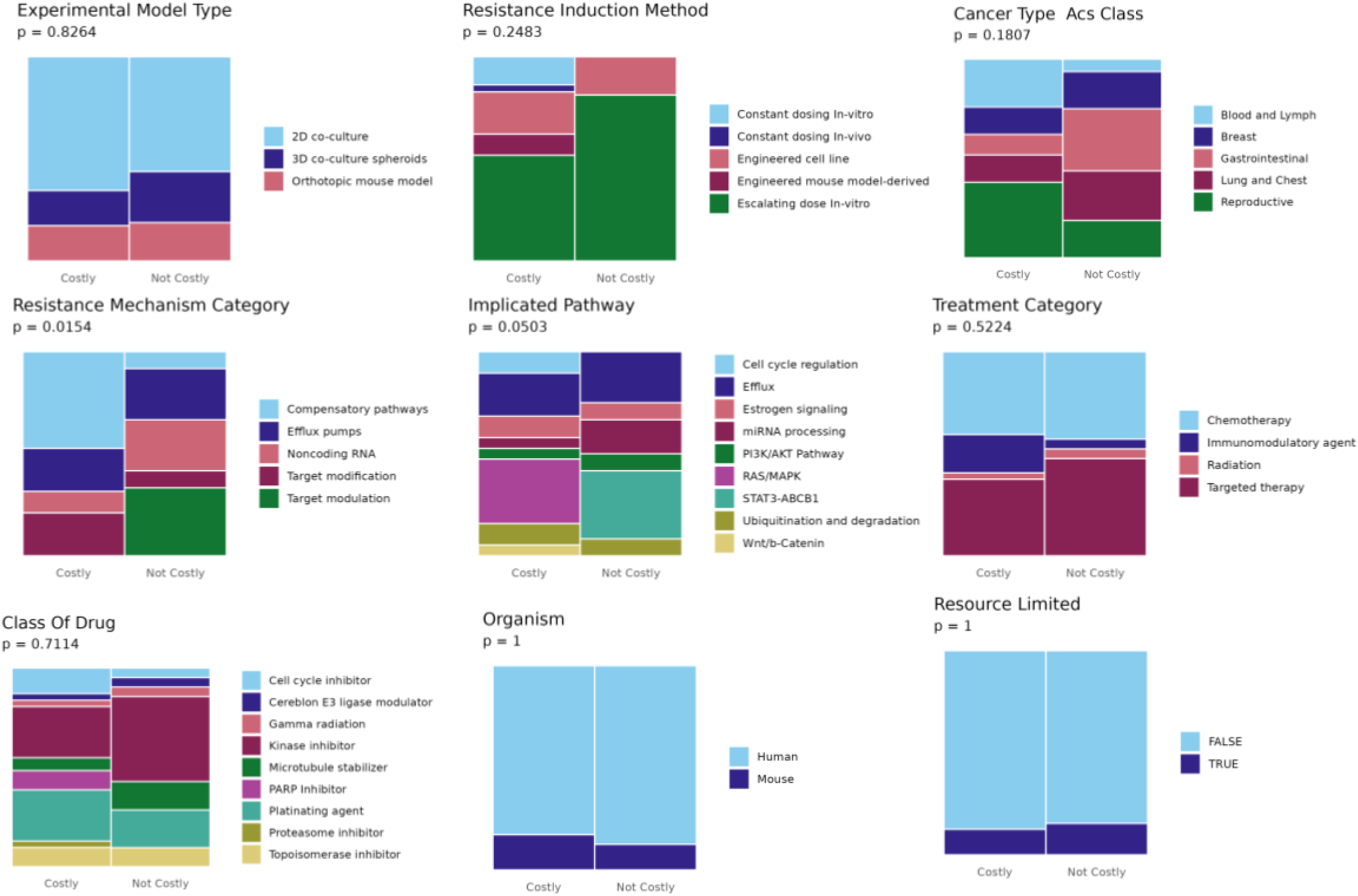
SUPPLEMENTAL FIGURE S2.

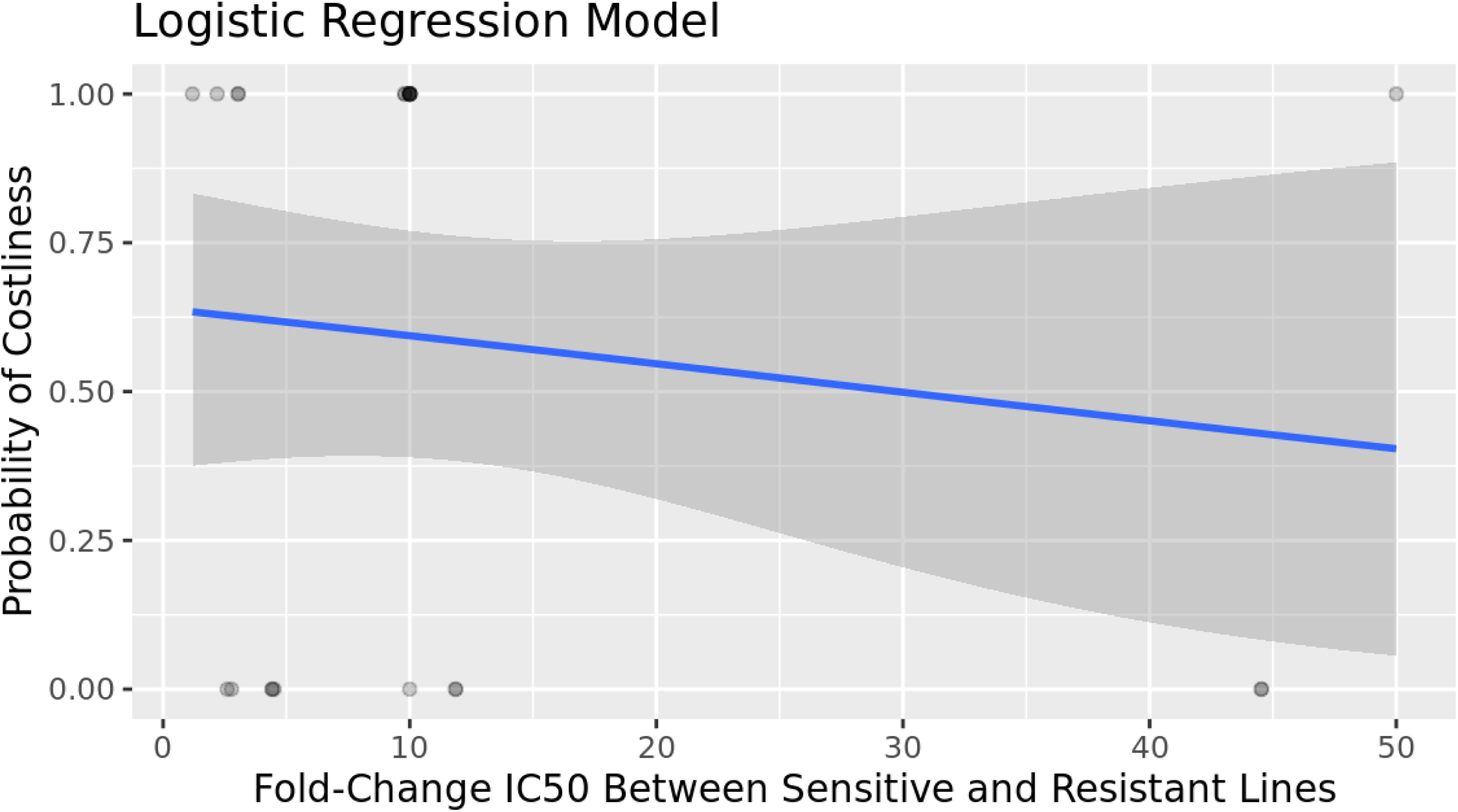
SUPPLEMENTAL FIGURE S3A.

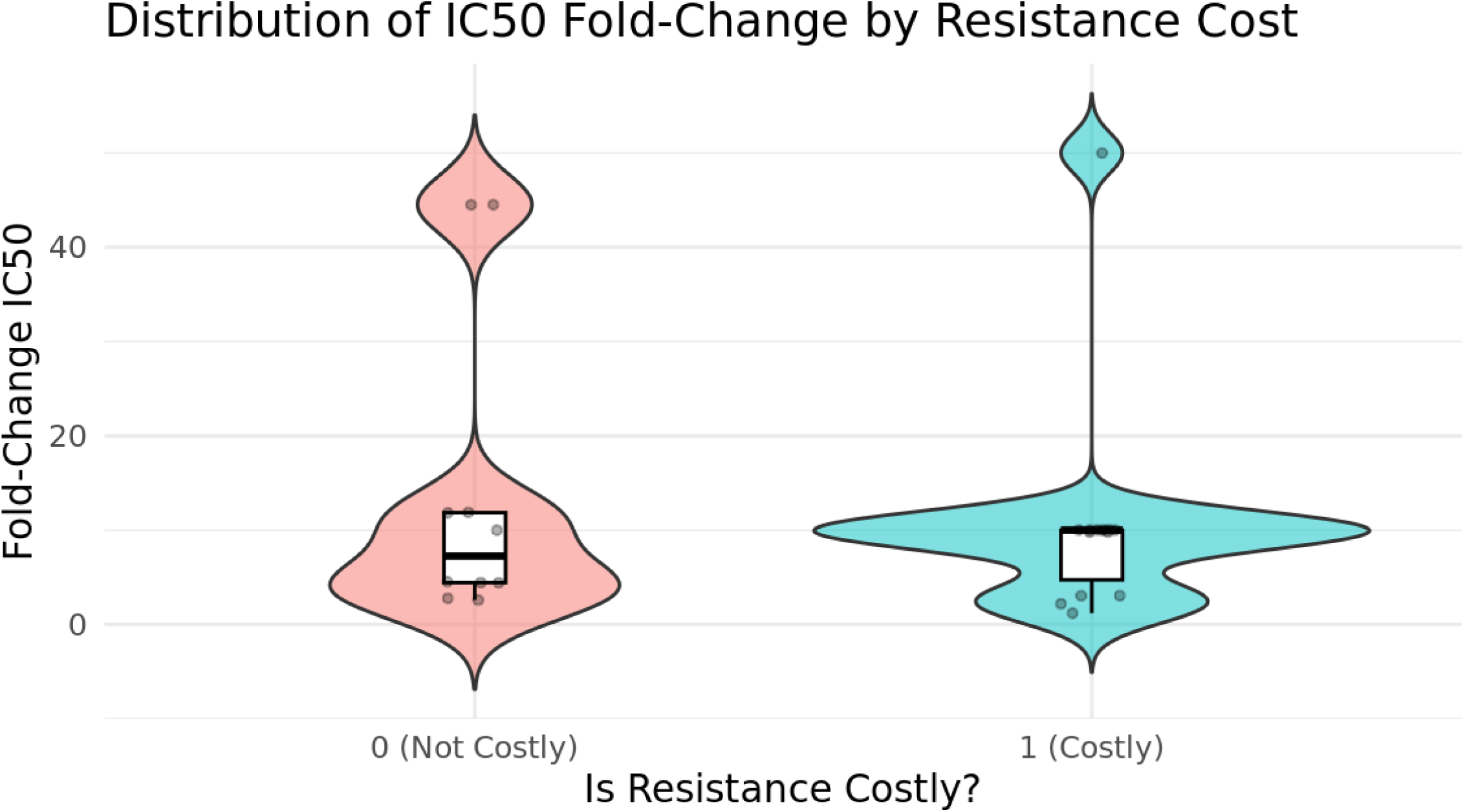
SUPPLEMENTAL FIGURE S3B.

