## Supplemental File 1 for "The Fitness Cost of Therapeutic Resistance in Cancer: A Systematic Review"

#### Supplemental Table S1. Search Strings and Dates Conducted

| Database | Date Searched | Query | No. Results |
|---|---|---|---|
| SciSpace | 9 September 2025 | “What is the fitness cost of drug resistance mutations in cancer cells and how do resistant clones compete with sensitive cells?” | 50 |
| Google Scholar | 3 October 2025 | cancer AND ("therapeutic resistance" OR "drug resistance") AND ("coculture" OR "coculture" OR "mixed culture" OR "competition assay" OR "clonal competition") AND "fitness" AND (trade*off OR *advantage OR select*) AND ("in vivo" or "in vitro") AND (evolution OR darwin OR selection) AND compete AND select)) -thesis -dissertation -fungal -antibiotic | 58 |
| Scopus | 3 October 2025 | ( TITLE-ABS-KEY ( ( "coculture" OR "coculture" OR "mixed culture" OR "competition assay" OR "clonal mixture" OR "heterogeneous culture" ) AND ( "sensitive" OR "drug-sensitive" OR "therapy-sensitive" OR "chemosensitive" ) AND ( "resistant" OR "drug-resistant" OR "therapy-resistant" OR "chemoresistant" ) AND ( "clonal competition" OR "clonal fitness" OR "cell competition" OR "competitive interactions" OR "cooperative interactions" OR "tumor evolution" OR "adaptive therapy" OR "evolutionary dynamics" ) AND ( cancer OR tumor OR tumour OR neoplasm OR carcinoma ) ) AND NOT ( fungal OR yeast OR bacteria OR antibiotic ) ) OR ( TITLE-ABS-KEY ( ( "drug-resistant" OR "therapy-resistant" OR "chemoresistant" ) AND ( "drug-sensitive" OR "therapy-sensitive" OR "chemosensitive" ) AND ( "cancer" OR "tumor" OR "tumour" OR "neoplasm" OR "carcinoma" ) AND ( "fitness" OR "growth rate" OR "competitive fitness" OR "cell competition" OR "proliferation rate" OR "competitive assay" OR "competition assay" OR "clonal competition" OR "evolutionary dynamics" OR "growth advantage" OR "fitness cost" ) AND ( "cell line" OR "coculture" OR "coculture" OR "in vitro" OR "culture experiment" ) ) AND NOT ( fungal OR yeast OR bacteria OR antibiotic ) ) OR ( TITLE-ABS-KEY ( competitive coculture cancer resistant ) ) OR ( ALL ( cancer AND ( "therapeutic resistance" OR "drug resistance" ) AND ( "coculture" OR coculture OR "mixed culture" OR "competition assay" OR "clonal competition" ) AND fitness AND ( "in vivo" OR "in vitro" ) AND ( evolution OR darwin OR selection ) ) AND NOT ALL ( fungal ) AND NOT ALL ( antibiotic ) AND NOT ALL ( bacterial ) ) AND ( EXCLUDE ( DOCTYPE , "re" ) ) | 85 |
| PubMed | 3 October 2025 | (("clonal competition"[tiab] OR "clonal fitness"[tiab] OR "fitness cost"[tiab] OR "competitive *advantage"[tiab] OR "growth cost"[tiab]) AND ("resistant clones"[tiab] OR "resistance mutations"[tiab] OR "drug resistance"[tiab] OR "therapeutic resistance"[tiab]) AND ("cancer"[tiab] OR "neoplasms"[mesh])) | 37 |
| SciSpace | 15 October 2025 | “Below is a list of 33 papers that have experiments where therapy-sensitive and therapy-resistant cancer cell lines are directly cocultured or otherwise mixed in a drug-free environment to evaluate fitness consequences of therapeutic resistance. Find related papers that meet this criteria without repeating any on the list. In-vivo or in-vitro models are acceptable.” | 0 |

#### Supplemental Figure S1. PRISMA2020 Chart and Sankey Diagram

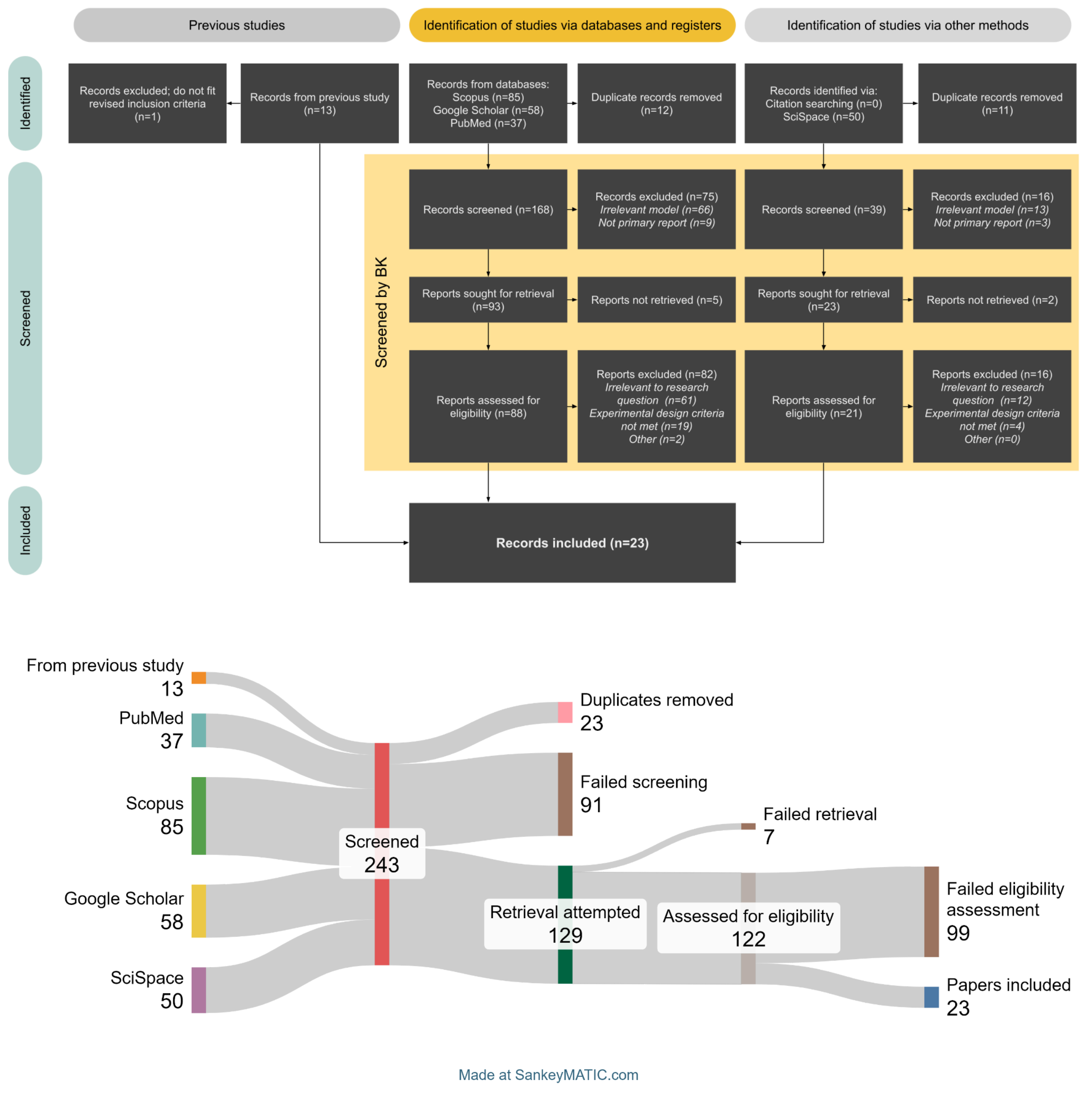

**Supplemental Figure S2. Outcome by Experimental Feature, Derived Populations Only**

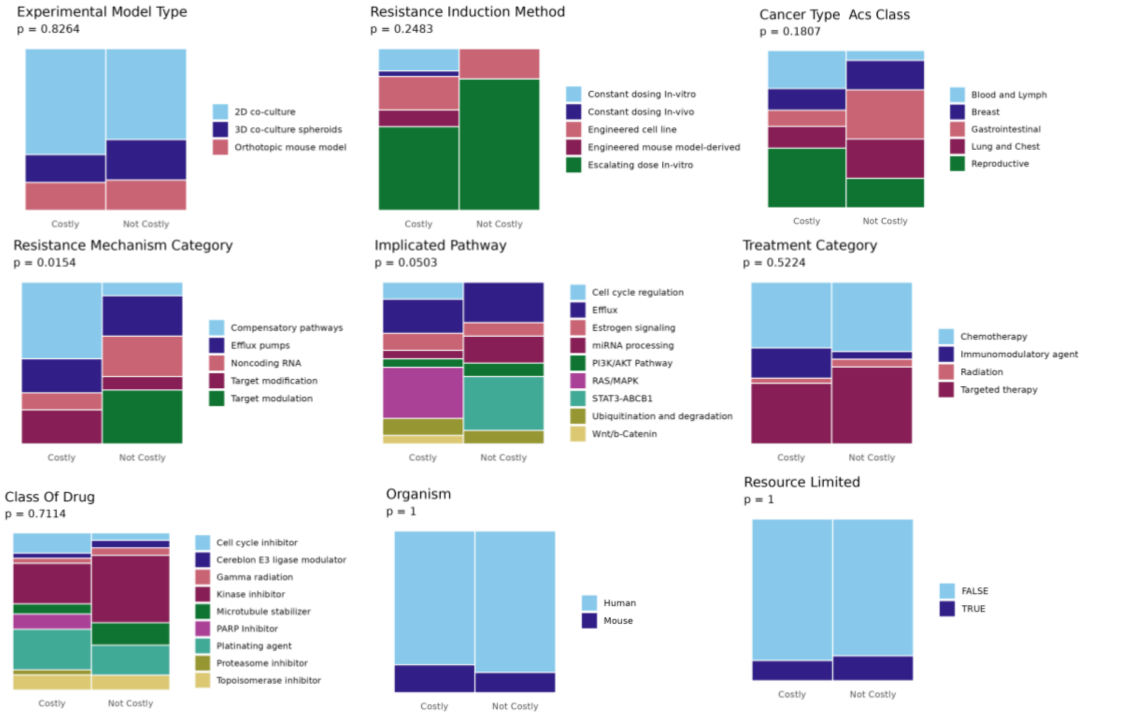


Mosaic plots depict distribution of experiment outcome among 9 experimental features. Experiments using non-isogenic cell lines (n=4) were excluded from this analysis. In order with associated column number from Supplemental Table S2: Experimental model type (Col. 5), resistance induction method (Col. 22), cancer type (Col. 11), resistance mechanism category (Col. 18), implicated pathway (Col. 20), treatment category (Col. 23), drug class (Col. 24), organism (Col. 10), resource limited *in vitro?* (Col. 9). Column height indicates proportion among group (Costly n=33, Not Costly n=16). Not Costly includes instances where resistant cells won in competition or no significant fitness gap was observed in competition.

**Supplemental Figure S3. Log Regression of Fold-Change IC50 Between Resistant and Sensitive Lines**

**A.**

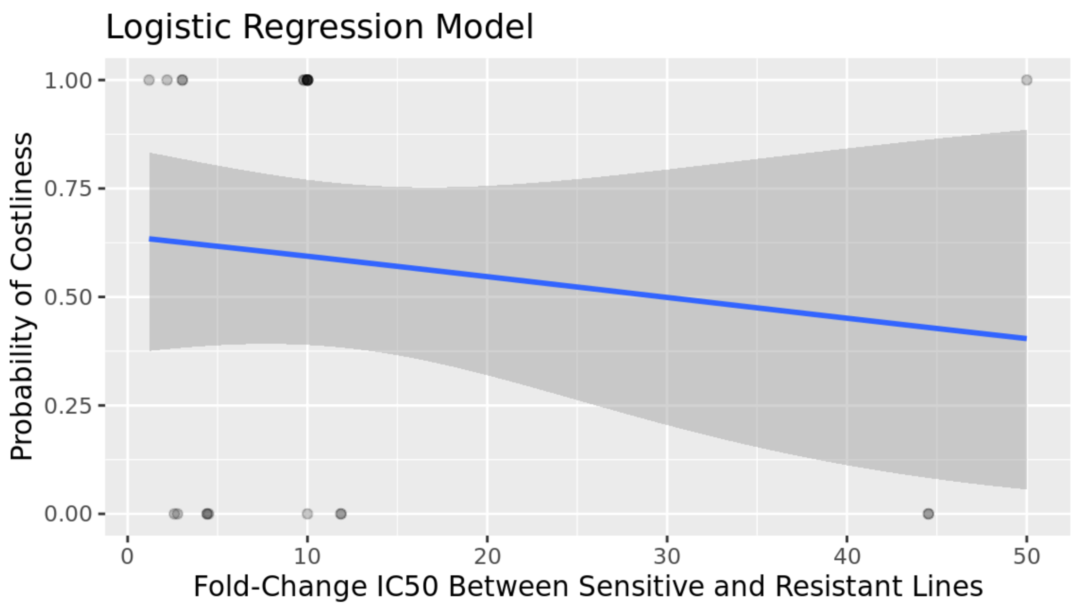


**B.**

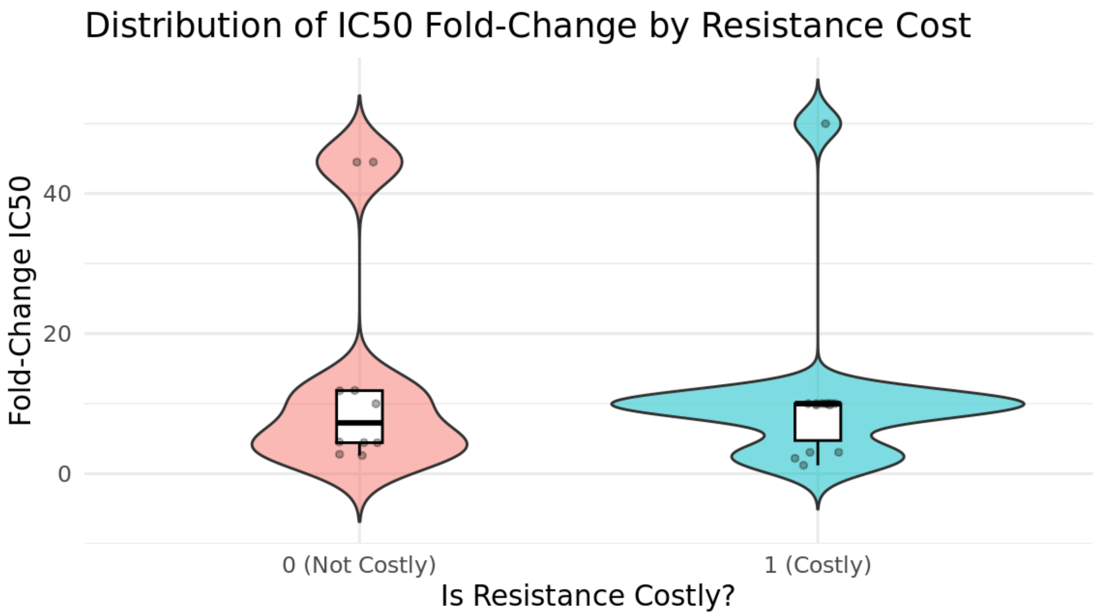


**A**. Logistic regression of fold-change IC50 between sensitive and resistant populations against whether resistance proved costly in competition (1 = costly, 0 = not costly). **B.** Violin plot showing distribution of fold-change IC50 between sensitive and resistant populations by outcome. No significant relationship was noted between magnitude of resistance and its costliness (p=0.535, Chi-Square Test).

**Supplemental Table S2. Data Collection Table Columns**

| # | Column Name | Description | Data Type | Options |
|---|---|---|---|---|
| 1 | Title | Title and citation of the publication | Nominal | *Unconstrained* |
| 2 | Experiment number (Exp. #) | If a publication contains multiple experiments; the experiment number | Numerical | 1-6 |
| 3 | Conclusion | The binary winner or loser status of resistant clones: did the resistant line win in coculture? | Nominal | Resistance does not confer a significant fitness difference<br>Resistance is beneficial<br>Resistance is costly |
| 4 | Modality | Describes if the experiment was conducted *in vivo* or *in vitro* | Nominal | *in vitro*<br>*in vivo* |
| 5 | Experimental model type | Describes the experimental model used | Nominal | 2D coculture<br>3D coculture spheroids<br>Orthotopic mouse model |
| 6 | Summary | A summary of the experiment | Nominal | *Unconstrained* |
| 7 | Measure of fitness | What metric was used to evaluate fitness? | Nominal | Population Growth Rate<br>Cellular proliferation rate<br>Cellular death rate |
| 8 | Fitness data collected by | What experimental techniques were used to measure fitness? | Nominal | *Unconstrained* |
| 9 | Resource limited? | Was the experiment conducted in resource-limited conditions? (*in vitro* experiments only) | Logical | TRUE<br>FALSE<br>NA |
| 10 | Cancer model | Describes the cancer cell line used and what species it is derived from | Nominal | *Unconstrained* |
| 11 | Cancer type ACS Class | Class of cancer by American Cancer Society classifications | Nominal | Blood and Lymph<br>Breast<br>Gastrointestinal<br>Lung and Chest<br>Reproductive |
| 12 | Resistant population derived? | If the resistant population was derived from the parental sensitive population | Logical | TRUE<br>FALSE<br>NA |
| 13 | Sensitive line | Name of the sensitive cell line used | Nominal | *Unconstrained* |
| 14 | Resistant line | Name of the resistant cell line used | Nominal | *Unconstrained* |
| 15 | Quantitative measure of resistance | How the study authors chose to measure resistance in their resistant cell line | Nominal | *Unconstrained* |
| 16 | Fold-change IC50 of resistant cell line | The fold-change in IC50 between sensitive and resistant cell lines, if provided | Numerical | *Unconstrained* |
| 17 | Admixture Ratio (Sens:Res) | The simplified ratio(s) of sensitive to resistant cells used in coculture, if provided | Numerical | *Unconstrained* |

| # | Column Name | Description | Data Type | Options |
|---|---|---|---|---|
| 18 | Resistance Mechanism category | The broad mechanism of resistance category, if provided | Nominal | Compensatory Pathways<br>Efflux Pumps<br>EMT<br>Metabolic Rewiring<br>Target Modulation<br>miRNA Dysregulation |
| 19 | Specific resistance characteristic | The specific mechanism of resistance, if identified | Nominal | *Unconstrained* |
| 20 | Implicated pathway | The molecular pathway involved in the given resistance mechanism | Nominal | Ubiquitination and degradation<br>Wnt/b-Catenin<br>RAS/MAPK<br>STAT3-ABCB1<br>Estrogen signaling<br>PI3K/AKT<br>miRNA processing<br>Cell cycle regulation<br>Efflux |
| 21 | Other resistance characteristics described | Other characteristics of the resistant cell described that do not fit the described mechanisms of resistance | Nominal | *Unconstrained* |
| 22 | Resistance induction method | The method by which resistant cell lines were generated for use in the study | Nominal | Engineered mouse model-derived<br>*in vitro* constant dosing<br>*in vitro* incremental dosing<br>*in vivo* constant dosing<br>Gene editing<br>Patient-derived |
| 23 | Treatment category | The class of treatment to which the resistant line is primarily tolerant; usually the class of drug to which resistance was induced, if applicable | Nominal | Chemotherapy<br>Immunomodulatory Agent<br>Multiple<br>Radiation<br>Targeted Therapy |
| 24 | Class of drug | The mechanism of action of the drug | Nominal | Cereblon E3 ligase modulator<br>Proteasome inhibitor<br>Platinating agent<br>Kinase inhibitor<br>Gamma radiation<br>Microtubule stabilizer<br>Topoisomerase inhibitor<br>Cell cycle inhibitor<br>Nucleic acid synthesis Inhibitor<br>PARP Inhibitor |
| 25 | Treatment name (trx name) | The specific name of the drug used, if applicable | Nominal | *Unconstrained* |
| 26 | Includes monoculture? | Does the study include a complementary experiment with the same parameters where cell lines are cultured separately? | Logical | TRUE<br>FALSE<br>NA |

| # | Column Name | Description | Data Type | Options |
| --- | --- | --- | --- | --- |
| 27 | Monoculture conclusion | What was the outcome of monoculture or homotypic culture experiments? How does the resistant growth rate compare to the sensitive growth rate when cultured separately? | Nominal | Resistance confers no significant intrinsic fitness difference<br>Resistance is intrinsically beneficial<br>Resistance is intrinsically costly |
| 28 | Monoculture summary | The summary of results from the monoculture experiment | Nominal | *Unconstrained* |

### Supplemental Table S3. Complete Table of Included Experiments

| Title | Exp. # | Conclusion | Modality | Experimental model type | Summary | Measure of fitness | Fitness data collected by | Resource limited? | Cancer model | Cancer Type (ACS Class) | Resistant population derived? | Sensitive line | Resistant line | Admixture ratio (Sens:Res) | Quantitative measure of resistance | Fold-change IC50 of resistant cells | Resistance mechanism category | Specific resistance mechanism | Implicated pathway | Other resistance characteristics described | Resistance induction method | Treatment category | Class of drug | Trx name | Includes monoculture? | Monoculture Conclusion | Monoculture Summary |
|---|---|---|---|---|---|---|---|---|---|---|---|---|---|---|---|---|---|---|---|---|---|---|---|---|---|---|---|
| IKZF1/3 and CRL4CRBN E3 ubiquitin ligase mutations and resistance to immunomodulatory drugs in multiple myeloma (Barrio et al., 2020) | 1 | Resistance is costly | *in vitro* | 2D coculture | CUL4B KO conferred lenalidomide resistance to multiple myeloma cells, but a growth penalty in coculture against parental cells | Population growth rate | Absolute number of cells by flow cytometry | FALSE | Human multiple myeloma | Blood and Lymph | TRUE | L363 CUL4B WT | L363 CUL4B KO | 1:1 | Clonal fraction under 2.5uM lenalidomide | NA | Target modification | CUL4B KO | Ubiquitination and degradation | NA | Engineered cell line | Immunomodulatory agent | Cereblon E3 ligase modulator | lenalidomide | FALSE | NA | NA |
| IKZF1/3 and CRL4CRBN E3 ubiquitin ligase mutations and resistance to immunomodulatory drugs in multiple myeloma (Barrio et al., 2020) | 2 | Resistance does not confer a significant fitness difference | *in vitro* | 2D coculture | IKZF1 A152T confers lenalidomide resistance to mulitple myeloma cells without significant fitness consequences in competition | Population growth rate | Absolute number of cells by flow cytometry | FALSE | Human multiple myeloma | Blood and Lymph | TRUE | L363 IKZF1 WT | L393 IKZF1 A152T | 9:1 | % Viability under 10uM lenalidomide | NA | Target modification | IKZF1 A12T | Ubiquitination and degradation | NA | Engineered cell line | Immunomodulatory agent | Cereblon E3 ligase modulator | lenalidomide | FALSE | NA | NA |
| Clonal competition assays identify fitness signatures in cancer progression and resistance in multiple myeloma (Haertle et al., 2024) | 1 | Resistance is costly | *in vitro* | 2D coculture | PSMB5 A20T conferred bortezomib resistance to multiple myeloma cells, but a growth penalty in coculture against parental cells | Population growth rate | Absolute number of cells by flow cytometry | FALSE | Human multiple myeloma | Blood and Lymph | TRUE | L363 PSMB5 WT | L363 PSMB5 A20T | 11:9 | RETRIEVAL ISSUE | NA | Target modification | PSMB5 A20T | Ubiquitination and degradation | NA | Engineered cell line | Targeted therapy | Proteasome inhibitor | bortezomib | FALSE | NA | NA |
| Adaptive therapy exploits fitness deficits in chemotherapy-resistant ovarian cancer to achieve long-term tumor control (Hockings et al., 2025) | 1 | Resistance confers benefit | *in vitro* | 2D coculture | coculture in high-resource conditions allowed for outcompetition of sensitive clones by cisplatin-resistant clones. Resistance induced **in-vitro.** | Population growth rate | Absolute number of cells by flow cytometry | FALSE | Human high-grade serious ovarian cancer | Reproductive | TRUE | OVCAR4 | Ov4Cis | 19:1, 17:3, 3:1, 1:1, 1:3, 3:17, 1:19 | Mean IC50 cisplatin | 10 | NA | NA | NA | ECM Remodeling gene expression changes | Escalating dose In-vitro | Chemotherapy | Platinating agent | cisplatin | TRUE | Resistance confers no significant intrinsic fitness difference in monoculture | Both cell lines displayed exponential growth in ad libitum conditions |
| Adaptive therapy exploits fitness deficits in chemotherapy-resistant ovarian cancer to achieve long-term tumor control (Hockings et al., 2025) | 2 | Resistance is costly | *in vitro* | 2D coculture | In resource-limited conditions, coculture resulted in outcompetition of cisplatin-resistant clones by sensitive clones. Greater rates of apoptosis and cell cycle arrest in resistant pop in coculture compared to monoculture. Resistance induced **in-vitro.** | Population growth rate | Absolute number of cells by flow cytometry | TRUE | Human high-grade serious ovarian cancer | Reproductive | TRUE | OVCAR4 | Ov4Cis | 17:3, 1:1, 3:17 | Mean IC50 cisplatin | 10 | NA | NA | NA | ECM Remodeling gene expression changes | Escalating dose In-vitro | Chemotherapy | Platinating agent | cisplatin | TRUE | Resistance confers no significant intrinsic fitness difference in monoculture | Both cell lines displayed reduced, but similar to one another, growth rates in limited-resource conditions |
| Adaptive therapy exploits fitness deficits in chemotherapy-resistant ovarian cancer to achieve long-term tumor control (Hockings et al., 2025) | 3 | Resistance is costly | *in vitro* | 2D coculture | In resource-limited conditions, coculture resulted in outcompetition of carboplatin-resistant clones by sensitive clones. Resistance induced **in-vitro.** | Population growth rate | Absolute number of cells by flow cytometry | TRUE | Human high-grade serious ovarian cancer | Reproductive | TRUE | OVCAR4 | Ov4Carbo | 17:3, 1:1, 3:17 | Mean IC50 carboplatin | 10 | NA | NA | NA | ECM Remodeling gene expression changes | Escalating dose In-vitro | Chemotherapy | Platinating agent | carboplatin | TRUE | Resistance confers no significant intrinsic fitness difference in monoculture | "mean subcutaneous tumor growth was comparable in OVCAR4 and Ov4Carbo cells although growth was variable between individual tumors" |
| Adaptive therapy exploits fitness deficits in chemotherapy-resistant ovarian cancer to achieve long-term tumor control (Hockings et al., 2025) | 4 | Resistance is costly | *in vitro* | 2D coculture | In resource-limited conditions, coculture resulted in outcompetition of carboplatin-resistant clones by sensitive clones. Resistance induced **in-vivo.** | Population growth rate | Absolute number of cells by flow cytometry | TRUE | Human high-grade serious ovarian cancer | Reproductive | TRUE | OVCAR4 | IVR01 | 17:3, 1:1, 3:17 | Mean IC50 carboplatin | 10 | NA | NA | NA | ECM Remodeling gene expression changes | Constant dosing In-vivo | Chemotherapy | Platinating agent | carboplatin | FALSE | NA | NA |
| Adaptive therapy exploits fitness deficits in chemotherapy-resistant ovarian cancer to achieve long-term tumor control (Hockings et al., 2025) | 5 | Resistance does not confer a significant fitness difference | *in vitro* | 2D coculture | In resource-limited conditions, coculture resulted in outcompetition of whichever clone was present in the lesser proportion. Resistance induced **in-vitro.** | Population growth rate | Absolute number of cells by flow cytometry | TRUE | Human high-grade serious ovarian cancer | Reproductive | TRUE | Cov318 | Cov-Cis | 17:3, 1:1, 3:17 | Mean IC50 cisplatin | 4.5 | NA | NA | NA | ECM Remodeling gene expression changes | Escalating dose In-vitro | Chemotherapy | Platinating agent | cisplatin | FALSE | NA | NA |
| Adaptive therapy exploits fitness deficits in chemotherapy-resistant ovarian cancer to achieve long-term tumor control (Hockings et al., 2025) | 6 | Resistance is costly | *in vivo* | Orthotopic mouse model | When mixed cultures of cisplatin-sensitive and -resistant clones were injected into mice, the proportion of resistant cells was decreased at 12 weeks. Regions of tumor derived from resistant cells had high Cas3 cleavage. | Population growth rate, Cellular proliferation rate, Cellular death rate | Amount of tag DNA by qPCR and approximate clonal contribution by IHC and pixel quantification | NA | Human high-grade serious ovarian cancer | Reproductive | TRUE | OVCAR4-GFP | Ov4Cis-RFP | 4:1, 1:1, 1:9 | Mean IC50 cisplatin | 10 | NA | NA | NA | ECM Remodeling gene expression changes | Escalating dose In-vitro | Chemotherapy | Platinating agent | cisplatin | FALSE | NA | NA |
| Evolution of Relapse-Proficient Subclones Constrained by Collateral Sensitivity to Oncogene Overdose in Wnt-Driven Mammary Cancer (Keller and Gunther, 2019) | 1 | Resistance is costly | *in vivo* | Orthotopic mouse model | Oncogene-addicted resistant lines cells were outcompeted by sensitive cells when targeted therapy was withheld in a Wnt-driven model of mouse mammary carcinoma | Population growth rate | Approximate clonal contribution by MRS-informed tumor segmentation | NA | Mouse mammary carcinoma | Breast | TRUE | P (iWnt/ApcM in parental mammary tumor) | R2 (iWnt/ApcM in mammary tumor relapse) | NA | NA | NA | Compensatory pathways | APCmmcr 1568*/1522*, APCmmcr 1568+T, APCmmcr 1522 A>T | Wnt/b-Catenin | NA | Engineered mouse model-derived | Targeted therapy | NA | NA | TRUE | Resistance confers no significant intrinsic fitness difference in monoculture | Homotypic injection of R2+R1 lines results in no significant selection on a line. |
| JAK2 inhibition mediates clonal selection of RAS pathway mutations in myeloproliferative neoplasms (Maslah et al., 2025) | 1 | Resistance is costly | *in vitro* | 2D coculture | Nras-mutant, ruxolitinib-resistant mouse bone marrow cells were outcompeted by sensitive cells in drug-free vehicle (DMSO) conditions | Population growth rate | Absolute number of cells by flow cytometry | FALSE | Mouse myeloproliferative neoplasm | Blood and Lymph | TRUE | NRAS WT Lin- bone marrow cells | NRAS G12D Lin- bone marrow cells | 1:1, 4:1 | Colony number under 1uM ruxolitinib | NA | Compensatory pathways | NRAS G12D | RAS/MAPK | NA | Engineered mouse model-derived | Immunomodulatory agent | Kinase inhibitor | ruxolitinib | FALSE | NA | NA |
| JAK2 inhibition mediates clonal selection of RAS pathway mutations in myeloproliferative neoplasms (Maslah et al., 2025) | 2 | Resistance is costly | *in vivo* | Orthotopic mouse model | Nras-mutant, ruxolitinib-resistant mouse bone marrow cells were outcompeted by empty casette, WT sensitive cells in drug-free vehicle (DMSO) conditions | Population growth rate | Absolute number of cells by flow cytometry | NA | Mouse myeloproliferative neoplasm | Blood and Lymph | TRUE | NRAS WT Lin- bone marrow cells | NRAS G12D Lin- bone marrow cells | 9:1 | Colony number under 1uM ruxolitinib | NA | Compensatory pathways | NRAS G12D | RAS/MAPK | NA | Engineered mouse model-derived | Immunomodulatory agent | Kinase inhibitor | ruxolitinib | FALSE | NA | NA |
| JAK2 inhibition mediates clonal selection of RAS pathway mutations in myeloproliferative neoplasms (Maslah et al., 2025) | 3 | Resistance is costly | *in vitro* | 2D coculture | Nras-mutant, ruxolitinib-resistant human bone marrow cells were outcompeted by sensitive cells in drug-free vehicle (DMSO) conditions | Population growth rate | Absolute number of cells by flow cytometry | FALSE | Human myeloproliferative neoplasm | Blood and Lymph | TRUE | HEL NRAS WT | HEL NRAS Q61K | 4:1 | Colony number under 1uM ruxolitinib | NA | Compensatory pathways | NRAS Q61K | RAS/MAPK | NA | Engineered cell line | Immunomodulatory agent | Kinase inhibitor | ruxolitinib | FALSE | NA | NA |
| JAK2 inhibition mediates clonal selection of RAS pathway mutations in myeloproliferative neoplasms (Maslah et al., 2025) | 4 | Resistance is costly | *in vitro* | 2D coculture | Nras-mutant, ruxolitinib-resistant human bone marrow cells were outcompeted by sensitive cells in drug-free vehicle (DMSO) conditions | Population growth rate | Absolute number of cells by flow cytometry | FALSE | Human myeloproliferative neoplasm | Blood and Lymph | TRUE | UKE-1 NRAS WT | UKE-1 NRAS Q61K | 4:1 | Colony number under 1uM ruxolitinib | NA | Compensatory pathways | NRAS Q61K | RAS/MAPK | NA | Engineered cell line | Immunomodulatory agent | Kinase inhibitor | ruxolitinib | FALSE | NA | NA |
| JAK2 inhibition mediates clonal selection of RAS pathway mutations in myeloproliferative neoplasms (Maslah et al., 2025) | 5 | Resistance is costly | *in vitro* | 2D coculture | Nras-mutant, ruxolitinib-resistant human bone marrow cells were outcompeted by sensitive cells in drug-free vehicle (DMSO) conditions | Population growth rate | Absolute number of cells by flow cytometry | FALSE | Humanized mouse pro-b cell | Blood and Lymph | TRUE | Ba/F3 NRAS WT | Ba/F3 NRAS Q61K | 4:1 | Colony number under 1uM ruxolitinib | NA | Compensatory pathways | NRAS Q61K | RAS/MAPK | NA | Engineered cell line | Immunomodulatory agent | Kinase inhibitor | ruxolitinib | FALSE | NA | NA |
| BNIP3-mediated mitophagy boosts the competitive growth of Lenvatinib-resistant cells via energy metabolism reprogramming in HCC (Wang et al., 2024) | 1 | Resistance confers benefit | *in vitro* | 2D coculture | Overexpression of RPTKs and glycolytic enhancement conferred both lenvatinib resistance and a growth advantage in vitro. | Population growth rate | Absolute number of cells by flow cytometry and high-content imaging | FALSE | Human hepatocellular carcinoma | Gastrointestinal | TRUE | CCHuh7m | CCHuh7R | 1:1 | Mean IC50 lenvatinib | 11.86 | Target modulation | RPTK hi | STAT3-ABCB1 | Glycolytic metabolism, enhanced mitophagy | Escalating dose In-vitro | Targeted therapy | Kinase inhibitor | lenvatinib | TRUE | Resistant cells have a significant growth benefit in monoculture | Sensitive Huh7 lines had decreased growth in competitive coculture compared to noncompetitive mono- or coculture. Further, sensitive cells from competitive coculture assays retained a growth disadvantage even in monoculture, though sensitive cells from noncompetitive cultures retained baseline proliferation rates. Monoculture of resistant line demonstrated increased proliferation rate compared to monoculture baseline proliferation for sensitive lines. |

| Title | Exp. # | Conclusion | Modality | Experimental model type | Summary | Measure of fitness | Fitness data collected by | Resource limited? | Cancer model | Cancer Type (ACS Class) | Resistant population derived? | Sensitive line | Resistant line | Admixture ratio (Sens:Res) | Quantitative measure of resistance | Fold-change IC50 of resistant cells | Resistance mechanism category | Specific resistance mechanism | Implicated pathway | Other resistance characteristics described | Resistance induction method | Treatment category | Class of drug | Trx name | Includes monoculture? | Monoculture Conclusion | Monoculture Summary |
|---|---|---|---|---|---|---|---|---|---|---|---|---|---|---|---|---|---|---|---|---|---|---|---|---|---|---|---|
| BNIP3-mediated mitophagy boosts the competitive growth of Lenvatinib-resistant cells via energy metabolism reprogramming in HCC (Wang et al., 2024) | 2 | Resistance confers benefit | *in vivo* | Orthotopic mouse model | RPTK-overexpressing lines with glycolytic enhancement conferred lenvatinib resistance and a competitive growth advantage in vivo | Population growth rate | Number of cells by tumor volume, approximate clonal contribution by fluorescence imaging and H&E stain, and absolute number of cells by flow cytometry and high-content imaging | NA | Human hepatocellular carcinoma | Gastrointestinal | TRUE | CCHuh7m | CCHuh7R | 1:1 | Mean IC50 lenvatinib | 11.86 | Target modulation | RPTK hi | STAT3-ABCB1 | Glycolytic metabolism, enhanced mitophagy | Escalating dose In-vitro | Targeted therapy | Kinase inhibitor | lenvatinib | TRUE | Resistant cells have a significant growth cost in monoculture | Huh7R cells alone yielded smaller tumors by volume. |
| BNIP3-mediated mitophagy boosts the competitive growth of Lenvatinib-resistant cells via energy metabolism reprogramming in HCC (Wang et al., 2024) | 3 | Resistance confers benefit | *in vitro* | 2D coculture | Overexpression of RPTKs and glycolytic enhancement conferred both lenvatinib resistance and a growth advantage in vitro | Population growth rate | Absolute number of cells by flow cytometry and high-content imaging | FALSE | Human hepatocellular carcinoma | Gastrointestinal | TRUE | CCPLC-PRF-5m | CCPLC-PRF-5R | 1:1 | Mean IC50 lenvatinib | 4.42 | Target modulation | RPTK hi | STAT3-ABCB1 | Glycolytic metabolism, enhanced mitophagy | Escalating dose In-vitro | Targeted therapy | Kinase inhibitor | lenvatinib | TRUE | Resistance confers no significant intrinsic fitness difference | mCherry-tagged sensitive HCC cell lines had decreased growth in competitive coculture compared to noncompetitive monoculture |
| BNIP3-mediated mitophagy boosts the competitive growth of Lenvatinib-resistant cells via energy metabolism reprogramming in HCC (Wang et al., 2024) | 4 | Resistance confers benefit | *in vivo* | Orthotopic mouse model | RPTK-overexpressing lines with glycolytic enhancement conferred lenvatinib resistance and a competitive growth advantage in vivo | Population growth rate | Number of cells by tumor volume, approximate clonal contribution by fluorescence imaging and H&E stain, and absolute number of cells by flow cytometry and high-content imaging | NA | Human hepatocellular carcinoma | Gastrointestinal | TRUE | CCPLC-PRF-5m | CCPLC-PRF-5R | 1:1 | Mean IC50 lenvatinib | 4.42 | Target modulation | RPTK hi | STAT3-ABCB1 | Glycolytic metabolism, enhanced mitophagy | Escalating dose In-vitro | Targeted therapy | Kinase inhibitor | lenvatinib | TRUE | Resistant cells have a significant growth cost in monoculture | PLC-PRF-5R cells alone yielded smaller tumors by volume. |
| Cell facilitation promotes growth and survival under drug pressure in breast cancer (Emond et al., 2023) | 1 | Resistance is costly | *in vitro* | 3D coculture spheroids | Growth rate of ribociclib-resistant CAMA-1 clones is reduced in coculture while sensitive clones see a growth advantage due to increased estradiol production by resistant clones. | Population growth rate | Approximate clonal contribution by fluorescence imaging | FALSE | Human ER+ breast cancer | Breast | TRUE | CAMA-1 ribociclib-sensitive | CAMA-1 ribociclib-resistant | 1:1 | Growth rate fold reduction under 400nM ribociclib relative to untreated conditions | NA | Compensatory pathways | Estradiol production hi | Estrogen signaling | NA | Escalating dose In-vitro | Targeted therapy | Cell cycle inhibitor | ribociclib | TRUE | Resistance confers no significant intrinsic fitness difference in monoculture | Sensitive cells in monoculture had a slower growth rate than when cocultured, suggesting a winner/loser dynamic. Resistant cells in monoculture have a similar growth rate to sensitive cells. |
| Cell facilitation promotes growth and survival under drug pressure in breast cancer (Emond et al., 2023) | 2 | Resistance is costly | *in vitro* | 3D coculture spheroids | Growth rate of ribociclib-resistant LY2 clones is reduced in coculture while sensitive clones see a growth advantage. | Population growth rate | Approximate clonal contribution by fluorescence imaging | FALSE | Human ER+ breast cancer | Breast | TRUE | LY2 ribociclib-sensitive | LY2 ribociclib-resistant | 4:1 | Growth rate under 400nM ribociclib relative to untreated conditions | NA | Compensatory pathways | Estradiol production hi | Estrogen signaling | NA | Escalating dose In-vitro | Targeted therapy | Cell cycle inhibitor | ribociclib | TRUE | Resistance confers no significant intrinsic fitness difference in monoculture | Both cell lines had similar growth rates in absence of therapy. |
| Cell facilitation promotes growth and survival under drug pressure in breast cancer (Emond et al., 2023) | 3 | Resistance confers benefit | *in vitro* | 3D coculture spheroids | Growth rate of ribociclib-resistant MCF7 clones is increased in coculture while sensitive clones see a growth disadvantage. | Population growth rate | Approximate clonal contribution by fluorescence imaging | FALSE | Human ER+ breast cancer | Breast | TRUE | MCF7 ribociclib-sensitive | MCF7 ribociclib-resistant | 9:1 | EC50 ribociclib | 2.77 | Compensatory pathways | Estradiol production hi | Estrogen signaling | NA | Escalating dose In-vitro | Targeted therapy | Cell cycle inhibitor | ribociclib | TRUE | Resistant cells have a significant growth benefit in monoculture | Resistant cell lines had highr growth rates than sensitive lines when cultured separately. |
| Reciprocal interactions between tumour cell populations enhance growth and reduce radiation sensitivity in prostate cancer (Paczkowski et al., 2021) | 1 | Resistance confers benefit | *in vitro* | 3D coculture spheroids | Growth rate of radiation-resistant PC3 cells is greater than that of rad-sensitive clones | Population growth rate, Cellular proliferation rate, Cellular death rate | Absolute number of cells by flow cytometry, proliferation rate by EdU assay, death rate by flow cytometry | FALSE | Human prostate cancer | Reproductive | TRUE | PC3 Parental | PC3 RR | 1:1 | Mean proliferation rate under 6Gy ionizing radiation | NA | Noncoding RNA | miR-95 upregulation targeting SGPP1 | PI3K/AKT Pathway | NA | Escalating dose In-vitro | Radiation | Gamma radiation | NA | TRUE | Resistant cells have a significant growth benefit in monoculture | Monoculture sensitive spheroids were smaller than monoculture resistant spheroids, and sensitive cells have lower growth rates in monoculture |
| Reciprocal interactions between tumour cell populations enhance growth and reduce radiation sensitivity in prostate cancer (Paczkowski et al., 2021) | 2 | Resistance is costly | *in vitro* | 3D coculture spheroids | Growth rate of radiation-resistant DU145 cells is reduced compared to sensitive lines | Population growth rate | Absolute number of cells by flow cytometry | FALSE | Human prostate cancer | Reproductive | TRUE | DU145 Parental | DU145 RR | 1:1 | Mean inactivation dose of ionizing radiation | 1.19 | Noncoding RNA | lncRNA UCA1 upregulation | PI3K/AKT Pathway | NA | Escalating dose In-vitro | Radiation | Gamma radiation | NA | TRUE | Resistant cells have a significant growth benefit in monoculture | Monoculture sensitive spheroids were smaller than monoculture resistant spheroids, and sensitive cells have lower growth rates in monoculture |
| Fibroblasts and alectinib switch the evolutionary games played by non-small cell lung cancer (Kaznatcheev et al., 2019) | 1 | Resistance confers benefit | *in vitro* | 2D coculture | In drug-free conditions, resistant clones have an increased growth rate compared to parental lines, despite the two having similar growth rates when cultured separately. | Population growth rate | Approximate clonal contribution by fluorescence imaging | FALSE | Human non-small cell lung cancer | Lung and Chest | TRUE | H3122 Parental | H3122 Resistant | 9:1, 4:1, 3:2, 2:3, 1:4, 1:9 | Growth rate under alectinib | NA | NA | NA | NA | NA | Escalating dose In-vitro | Targeted therapy | Kinase inhibitor | alectinib | TRUE | Resistance confers no significant intrinsic fitness difference in monoculture | Resistant and parental lines have similar growth rates in monoculture. |
| Cooperative adaptation to therapy (CAT) confers resistance in heterogeneous non-small cell lung cancer (Craig et. al, 2019) | 1 | Resistance is costly | *in vitro* | 3D coculture spheroids | In coculture, resistant Dicer1 mutant cells have a slower growth rate than sensitie parental cells | Population growth rate | Approximate clonal contribution by fluorescence imaging and absolute number of cells by flow cytometry | FALSE | Mouse non-small cell lung cancer | Lung and Chest | TRUE | Dicer1 WT | Dicer1 M1 mutant | 9:1, 1:1, 1:9 | Number of cells under 96h multidrug exposure compared to vehicle control | NA | Noncoding RNA | Various unspecified Dicer1 mutations | miRNA processing | NA | Engineered cell line | Targeted therapy, Chemotherapy, Targeted therapy | Kinase inhibitor, Microtubule stabilizer, Kinase inhibitor | afatinib, docetaxel, bortezomib | TRUE | Resistance confers no significant intrinsic fitness difference in monoculture | Resistant and parental lines have similar growth rates in monoculture. |
| Cooperative adaptation to therapy (CAT) confers resistance in heterogeneous non-small cell lung cancer (Craig et. al, 2019) | 2 | Resistance confers benefit | *in vitro* | 3D coculture spheroids | In coculture, Dicer1 mutants have a growth advantage over sensitive clones in absence of drug | Population growth rate | Approximate clonal contribution by fluorescence imaging and absolute number of cells by flow cytometry | FALSE | Mouse non-small cell lung cancer | Lung and Chest | TRUE | Dicer1 WT | Dicer1 M2 mutant | 1:1 | Number of cells under 96h multidrug exposure compared to vehicle control | NA | Noncoding RNA | Various unspecified Dicer1 mutations | miRNA processing | NA | Engineered cell line | Targeted therapy, Chemotherapy, Targeted therapy | Kinase inhibitor, Microtubule stabilizer, Kinase inhibitor | afatinib, docetaxel, bortezomib | TRUE | Resistance confers no significant intrinsic fitness difference in monoculture | Resistant and parental lines have similar growth rates in monoculture. |
| Cooperative adaptation to therapy (CAT) confers resistance in heterogeneous non-small cell lung cancer (Craig et. al, 2019) | 3 | Resistance confers benefit | *in vitro* | 3D coculture spheroids | In coculture, Dicer1 mutants have a growth advantage over sensitive clones in absence of drug | Population growth rate | Approximate clonal contribution by fluorescence imaging and absolute number of cells by flow cytometry | FALSE | Mouse non-small cell lung cancer | Lung and Chest | TRUE | Dicer1 WT | Dicer1 M3 mutant | 1:1 | Number of cells under 96h multidrug exposure compared to vehicle control | NA | Noncoding RNA | Various unspecified Dicer1 mutations | miRNA processing | NA | Engineered cell line | Targeted therapy, Chemotherapy, Targeted therapy | Kinase inhibitor, Microtubule stabilizer, Kinase inhibitor | afatinib, docetaxel, bortezomib | TRUE | Resistance confers no significant intrinsic fitness difference in monoculture | Resistant and parental lines have similar growth rates in monoculture. |
| Spatial heterogeneity and evolutionary dynamics modulate time to recurrence in continuous and adaptive cancer therapies (Gallaher et al., 2018) | 1 | Resistance is costly | *in vitro* | 2D coculture | In coculture, doxorubicin-resistant MCF7 clones lose against sensitive MCF7 clones | Population growth rate | Absolute cell number by flow cytometry and fluorescence imaging | FALSE | Human ER+ breast cancer | Breast | TRUE | MCF7 | MCF7Dox | 1:1 | NA | NA | Efflux pumps | Pgp hi | Efflux | NA | Constant dosing In-vitro | Chemotherapy | Topoisomerase inhibitor | doxorubicin | TRUE | Resistant cells have a significant growth cost in monoculture | Resistant lines had a much slower growth rate than sensitive lines in monoculture |

| Title | Exp. # | Conclusion | Modality | Experimental model type | Summary | Measure of fitness | Fitness data collected by | Resource limited? | Cancer model | Cancer Type (ACS Class) | Resistant population derived? | Sensitive line | Resistant line | Admixture ratio (Sens:Res) | Quantitative measure of resistance | Fold-change IC50 of resistant cells | Resistance mechanism category | Specific resistance mechanism | Implicated pathway | Other resistance characteristics described | Resistance induction method | Treatment category | Class of drug | Trx name | Includes monoculture? | Monoculture Conclusion | Monoculture Summary |
|---|---|---|---|---|---|---|---|---|---|---|---|---|---|---|---|---|---|---|---|---|---|---|---|---|---|---|---|
| A Strategy to Delay the Development of Cisplatin Resistance by Maintaining a Certain Amount of Cisplatin-Sensitive Cells (Duan et al., 2017) | 1 | Resistance is costly | *in vivo* | Orthotopic mouse model | 1:1 HeLa:HeLa/ddp tumors displayed poor growth in vivo compared to sensitive-only tumors. No apparent growth rate change was observed for sensitive HeLa cells in mixed-culture tumors compared to sensitive-only tumors, but resistant cells were almost completely absent at 40 days post transplantation. | Population growth rate, Cellular proliferation rate, Cellular death rate | Tumor volume, cell cycle status by Ki67, and apoptosis rate by TUNEL staining | NA | Human cervical adenocarcinoma | Reproductive | TRUE | HeLa-RFP | HeLa/ddp | 1:1 | Mean IC50 cisplatin | 10 | NA | NA | NA | Glycolytic metabolism | Escalating dose In-vitro | Chemotherapy | Platinating agent | cisplatin | TRUE | Resistant cells have a significant growth cost in monoculture | Sensitive-only tumors grew large, while resistant-only tumors remained at a small volume despite the same number of tumors cells being injected. |
| Spatial competition constrains resistance to targeted cancer therapy (Bacevic et al., 2019) | 1 | Resistance is costly | *in vitro* | 2D coculture | In coculture, CDK inhibitor-resistant HCT116 cells experienced a growth disadvantage compared to sensitive clones | Population growth rate | Approximate clonal contribution by fluorescence imaging and absolute number of cells by flow cytometry | FALSE | Human colorectal cancer | Gastrointestinal | TRUE | GFP+CDKi-sensitive HCT116 | mCherry+R50 HCT116 | 1:1, 9:1, 99:1 | Cell count under 50uM NU6102 | NA | Target modification | CDK2 mutation | Cell cycle regulation | NA | Constant dosing In-vitro | Targeted therapy | Cell cycle inhibitor | NU6102 | TRUE | Resistant cells have a significant growth cost in monoculture | WT monocultures have a higher growth rate than res., suggesting intrinsic fitness penalty |
| Spatial competition constrains resistance to targeted cancer therapy (Bacevic et al., 2019) | 2 | Resistance is costly | *in vitro* | 3D coculture spheroids | In 3D coculture, CDK inhibitor-resistant HCT116 cells experienced a growth disadvantage compared to sensitive clones | Population growth rate | Approximate clonal contribution by fluorescence imaging and absolute number of cells by flow cytometry | FALSE | Human colorectal cancer | Gastrointestinal | TRUE | GFP+CDKi-sensitive HCT116 | mCherry+R50 HCT116 | 99:1 | Cell count under 50uM NU6102 | NA | Target modification | CDK2 mutation | Cell cycle regulation | NA | Constant dosing In-vitro | Targeted therapy | Cell cycle inhibitor | NU6102 | FALSE | NA | NA |
| Evolutionary approaches to prolong progression-free survival in breast cancer (Silva et al., 2012) | 1 | Resistance confers benefit | *in vitro* | 2D coculture | In high-glucose conditions, MCF7/Dox mutants had a slightly shorter doubling time than parental cells. | Population growth rate | Approximate cell number by crystal violet staining and fluorescence imaging | FALSE | Human ER+ breast cancer | Breast | TRUE | MCF7 | MCF7/Dox | 2:1 | Mean IC50 doxorubicin | 44.53 | Efflux pumps | Pgp hi | Efflux | NA | Escalating dose In-vitro | Chemotherapy | Topoisomerase inhibitor | doxorubicin | TRUE | Resistance confers no significant intrinsic fitness difference in monoculture | Sensitive and resistant lines had similr growth rates in hi-glucose conditions. |
| Evolutionary approaches to prolong progression-free survival in breast cancer (Silva et al., 2012) | 2 | Resistance confers benefit | *in vitro* | 2D coculture | In low-glucose conditions, MCF7/Dox mutants had a slightly shorter doubling time than parental cells. | Population growth rate | Approximate cell number by crystal violet staining and fluorescence imaging | TRUE | Human ER+ breast cancer | Breast | TRUE | MCF7 | MCF7/Dox | 2:1 | Mean IC50 doxorubicin | 44.53 | Efflux pumps | Pgp hi | Efflux | NA | Escalating dose In-vitro | Chemotherapy | Topoisomerase inhibitor | doxorubicin | TRUE | Resistant cells have a significant growth cost in monoculture | Sensitive lines had higher growth rates in low-glucose conditions compared to resistant lines, which had a sharp dropoff after 72h. |
| Adriamycin-resistant cells are significantly less fit than adriamycin-sensitive cells in cervical cancer. (Qi et al., 2021) | 1 | Resistance is costly | *in vivo* | Orthotopic mouse model | Mixed tumors consisting of 1:1 initial seeding ratio grew as large as fully parental tumors and consisted of nearly entirely parental cells by endpoint | Population growth rate | Tumor volume and approximate clonal contribution by fluorescence imaging | NA | Human cervical adenocarcinoma | Reproductive | TRUE | HeLa-RFP | HeLa/ADR | 1:1 | Mean IC50 adriamycin | 10 | NA | NA | NA | NA | Escalating dose In-vitro | Chemotherapy | Topoisomerase inhibitor | doxorubicin | TRUE | Resistant cells have a significant growth cost in monoculture | Resistant-only tumors were far smaller than parental tumors |
| Dynamic Phenotypic Switching and Group Behavior Help Non-Small Cell Lung Cancer Cells Evade Chemotherapy. (Nam et al., 2022) | 1 | Resistance is costly | *in vitro* | 2D coculture | both 12-hour and 3-week cocultures of H23 and H2009 cells resulted in a growth advantage of sens over res due to a secreted factor from sensitive cells | Population growth rate | Approximate clonal contribution by continuous fluorescence imaging | FALSE | Human non-small cell lung cancer | Lung and Chest | FALSE | S (H23) | T (H2009) | 8:1, 2:1, 1:1, 1:2, 1:8 | Mean IC50 cisplatin | NA | NA | NA | NA | NA | Patient-derived | Chemotherapy | Platinating agent | cisplatin | FALSE | NA | NA |
| Paracrine behaviors arbitrate parasite-like interactions between tumor subclones (Noble et al., 2021) | 1 | Resistance is costly | *in vitro* | 2D coculture | "Two cell lines derived from a single mouse mammary carcinoma – 168 and 4T07 cells – have similar growth rates when cultured individually, yet the 4T07 clone displays a dominant phenotype when grown together" | Population growth rate, Cellular proliferation rate, Cellular death rate | Absolute number of cells and death rate by flow cytometry | FALSE | Mouse mammary carcinoma | Breast | FALSE | 4T07 | 168FARN | 1:4, 1:3, 3:1, and 4:1 | RETRIEVAL ISSUE | NA | NA | NA | NA | NA | Escalating dose In-vitro | Chemotherapy | Nucleic acid synthesis Inhibitor | 2,6-diamino purine | TRUE | Resistance confers no significant intrinsic fitness difference in monoculture | Similar growth rates in monoculture |
| E2F1 mediates competition, proliferation and response to cisplatin in cohabitating resistant and sensitive ovarian cancer cells (Valdivia et al., 2024) | 1 | Resistance is costly | *in vitro* | 2D coculture | coculture of OVCAR5 and resistant derivatives resulted in a growth advantage of sensitive cells at a penalty to res. | Population growth rate | Absolute number of cells by flow cytometry | FALSE | Human ovarian cancer | Reproductive | TRUE | OVCAR5 | OVCAR5 CisR | 1:1, 1:2, 1:5, 1:7 | Mean IC50 cisplatin | 2.19 | NA | NA | NA | NA | Escalating dose In-vitro | Chemotherapy | Platinating agent | cisplatin | TRUE | Resistant cells have a significant growth cost in monoculture | Sens cells grow faster in coculture than in monoculture, and res cells grow slower |
| E2F1 mediates competition, proliferation and response to cisplatin in cohabitating resistant and sensitive ovarian cancer cells (Valdivia et al., 2024) | 2 | Resistance is costly | *in vitro* | 2D coculture | coculture of PE01 and resistant PE04 cells resulted in a growth advantage of sensitive cells at a penalty to res. | Population growth rate | Absolute number of cells by flow cytometry | FALSE | Human ovarian cancer | Reproductive | FALSE | PE01 | PE04 | 1:1, 1:2, 1:5, 1:7 | Mean IC50 cisplatin | 3.04 | NA | NA | NA | NA | Patient-derived | Chemotherapy | Platinating agent | cisplatin | TRUE | Resistant cells have a significant growth cost in monoculture | Sens cells grow faster in coculture than in monoculture, and res cells grow slower |
| E2F1 mediates competition, proliferation and response to cisplatin in cohabitating resistant and sensitive ovarian cancer cells (Valdivia et al., 2024) | 3 | Resistance is costly | *in vitro* | 2D coculture | Long-term coculture of PE01 and resistant PE04 cells resulted in a growth advantage of sensitive cells at a penalty to res. | Population growth rate | Approximate clonal contribution by fluorescence imaging | FALSE | Human ovarian cancer | Reproductive | FALSE | PE01 | PE04 | 1:2 | Mean IC50 cisplatin | 3.04 | NA | NA | NA | NA | Patient-derived | Chemotherapy | Platinating agent | cisplatin | FALSE | NA | NA |
| Metronomic Chemotherapy Modulates Clonal Interactions to Prevent Drug Resistance in Non-Small Cell Lung Cancer. (Bondarenko et al., 2021) | 1 | Resistance is costly | *in vitro* | 2D coculture | Drug-sensitive A549 clones inhibit the proliferation of the drug-resistant A549/EpoB40 clones | Population growth rate | Approximate clonal contribution by fluorescence imaging | FALSE | Human non-small cell lung cancer | Lung and Chest | TRUE | A549 | A549/EpoB40 | 23:5 | Mean IC50 cisplatin (approx), Mean IC50 patupilone (approx) | 8, 100 | NA | NA | NA | NA | Escalating dose In-vitro | Chemotherapy, Chemotherapy | Platinating agent Microtubule stabilizer | cisplatin, patupilone | TRUE | Resistant cells have a significant growth cost in monoculture | Res cells grow slower in coculture compared to monoculture |
| Metronomic Chemotherapy Modulates Clonal Interactions to Prevent Drug Resistance in Non-Small Cell Lung Cancer. (Bondarenko et al., 2021) | 2 | Resistance is costly | *in vitro* | 2D coculture | Drug-sensitive HT29 clones inhibit the proliferation of the drug-resistant HT29/Rox1 clones | Population growth rate | Approximate clonal contribution by fluorescence imaging | FALSE | Human colorectal cancer | Gastrointestinal | TRUE | HT29 | HT29/Rox1 | 23:5 | Mean IC50 oxaliplatin (approx) | 10 | NA | NA | NA | NA | Escalating dose In-vitro | Chemotherapy | Platinating agent | oxaliplatin | FALSE | NA | NA |
| Metronomic Chemotherapy Modulates Clonal Interactions to Prevent Drug Resistance in Non-Small Cell Lung Cancer. (Bondarenko et al., 2021) | 3 | Resistance is costly | *in vitro* | 2D coculture | Drug-sensitive A549 clones inhibit the proliferation of the drug-resistant A549/EVP16 clones | Population growth rate | Approximate clonal contribution by fluorescence imaging | FALSE | Human non-small cell lung cancer | Lung and Chest | TRUE | A549 | A549/VP16 | 23:5 | Mean IC50 etoposide (approx) | 50 | NA | NA | NA | NA | Escalating dose In-vitro | Chemotherapy | Topoisomerase inhibitor | etoposide | FALSE | NA | NA |
| Metronomic Chemotherapy Modulates Clonal Interactions to Prevent Drug Resistance in Non-Small Cell Lung Cancer. (Bondarenko et al., 2021) | 4 | Resistance does not confer a significant fitness difference | *in vivo* | Orthotopic mouse model | "tumor cell composition did not change over time with a mixture of drug-sensitive A549-mtDsRed and drug-resistant A549/EpoB40-GFP" | Population growth rate | Absolute number of cells by flow cytometry | NA | Human non-small cell lung cancer | Lung and Chest | TRUE | A549 | A549/EpoB40 | 7:3 | Mean IC50 cisplatin (approx), Mean IC50 patupilone (approx) | 8, 100 | NA | NA | NA | NA | Escalating dose In-vitro | Chemotherapy, Chemotherapy | Platinating agent Microtubule stabilizer | cisplatin, patupilone | FALSE | NA | NA |
| HSP90a promotes the resistance to oxaliplatin in HCC through regulating IDH1-induced cell competition (Wang et al., 2024) | 1 | Resistance confers benefit | *in vitro* | 2D coculture | Oxaliplatin resistance mediated by lipid metabolism allows resistant cells to win in coculture | Population growth rate | Absolute cell number by flow cytometry and fluorescence imaging | FALSE | Human hepatocellular carcinoma | Gastrointestinal | TRUE | MHCC97H-GFP | MHCC97H-OXR | 1:1 | Mean IC50 oxaliplatin (approx) | 2.6 | Efflux pumps | MDR1 hi | Efflux | Increased lipid metabolism | Escalating dose In-vitro | Chemotherapy | Platinating agent | oxaliplatin | FALSE | NA | NA |

| Title | Exp. # | Conclusion | Modality | Experimental model type | Summary | Measure of fitness | Fitness data collected by | Resource limited? | Cancer model | Cancer Type (ACS Class) | Resistant population derived? | Sensitive line | Resistant line | Admixture ratio (Sens:Res) | Quantitative measure of resistance | Fold-change IC50 of resistant cells | Resistance mechanism category | Specific resistance mechanism | Implicated pathway | Other resistance characteristics described | Resistance induction method | Treatment category | Class of drug | Trx name | Includes monoculture? | Monoculture Conclusion | Monoculture Summary |
|---|---|---|---|---|---|---|---|---|---|---|---|---|---|---|---|---|---|---|---|---|---|---|---|---|---|---|---|
| Establishment and Molecular Characterization of an In Vitro Model for PARPi-Resistant Ovarian Cancer (Klotz et al., 2023) | 1 | Resistance is costly | *in vitro* | 2D coculture | "In the coculture setting, we observed that Olres-UWB cells had a clear competitive disadvantage compared to PARPi-sensitive UWB cells" | Population growth rate | Absolute number of cells by flow cytometry | FALSE | Human ovarian cancer | Reproductive | TRUE | tdTomato-UWB1.289 | eGFP-Olres-UWB1.289 | 1:1 | Mean IC50 olaparib | 9.8 | Efflux pumps | Pgp hi | Efflux | Epithelial-mesenchymal transition | Escalating dose In-vitro | Targeted therapy | PARP Inhibitor | olaparib | TRUE | Resistance confers no significant intrinsic fitness difference in monoculture | "There was no significant difference in basal proliferation between PARPi-resistant Olres-UWB vs. parental PARPi-sensitive cells in monocultures" |
| Establishment and Molecular Characterization of an In Vitro Model for PARPi-Resistant Ovarian Cancer (Klotz et al., 2023) | 2 | Resistance is costly | *in vitro* | 2D coculture | "We observed similar clonal dynamics in this model, including a competitive disadvantage of Olres-UWB+BRCA1 under drug-free conditions" | Population growth rate | Absolute number of cells by flow cytometry | FALSE | Human ovarian cancer | Reproductive | TRUE | tdTomato-UWB1.289+BRCA1 | eGFP-Olres-UWB1.289+BRCA1 | 1:1 | Mean IC50 olaparib | 9.8 | Efflux pumps | Pgp hi | Efflux | Epithelial-mesenchymal transition | Escalating dose In-vitro | Targeted therapy | PARP Inhibitor | olaparib | TRUE | Resistant cells have a significant growth cost in monoculture | Resistant lines slightly lower growth rate in monoculture |
| Evolutionary dynamics of cancer multidrug resistance in response to olaparib and photodynamic therapy (Baglo et al., 2021) | 1 | Resistance is costly | *in vitro* | 2D coculture | "The sensitive variant OVCAR-8-DsRed2 quickly begins dominating the population within days of culture and is at around 95%" | Population growth rate | Approximate clonal contribution by fluorescence imaging and absolute number of cells by flow cytometry | FALSE | Human ovarian cancer | Reproductive | TRUE | OVCAR-8-DsRed2 | NCI/ADR-RES-EGFP | 1:1 | Cell count under 25uM olaparib | NA | Efflux pumps | Pgp hi | Efflux | NA | Escalating dose In-vitro | Targeted therapy | PARP Inhibitor | olaparib | TRUE | Resistant cells have a significant growth cost in monoculture | Resistant lines slightly lower growth rate in monoculture |
| Measuring competitive exclusion in non–small cell lung cancer (Farrokhian et al., 2022) | 1 | Resistance is costly | *in vitro* | 2D coculture | " Ecological interactions alter but do not ameliorate the resistant clones fitness cost and result in competitive exclusion of the resistant strain in DMSO." | Population growth rate | Approximate clonal contribution by fluorescence imaging | FALSE | Human lung adenocarcinoma | Lung and Chest | TRUE | Sensitive PC9 | Resistant PC9 | Varied, from 9:1 to 1:9 | Growth rate under gefitinib | NA | Compensatory pathways | KRAS G12D | RAS/MAPK | EGFR downregulation | Constant dosing In-vitro | Targeted therapy | Kinase inhibitor | gefitinib | TRUE | Resistant cells have a significant growth cost in monoculture | Resistant lines have a slower growth rate than sensitive lines in monoculture |
